# APOSM: Pairwise preference learning improves generative small-molecule design

**DOI:** 10.64898/2026.06.06.730554

**Authors:** Marcus W. Dreisler, Richard Michael, Nikos Hatzakis, Wouter Boomsma

## Abstract

Small-molecule lead refinement is constrained by the cost of synthesizing and assaying candidates, making the surrogate models that prioritize compounds for experimental testing central to the design process. The reliability of such surrogates is limited by the noise and sparsity of screening measurements. We show that training the surrogate on pairwise comparisons between candidate molecules, rather than on absolute predicted scores, yields a substantially more reliable signal for active candidate selection in this regime. We develop APOSM, an active-learning algorithm that combines a fragment-based generator, a pairwise message-passing graph neural network surrogate, and probabilistic ranking inside a batched acquisition loop. On the Practical Molecular Optimization benchmark and a GPCR ligand rediscovery task, APOSM improves target attainment and sampling efficiency over unguided fragment-based optimization, the Graph-GA genetic algorithm, and a pointwise-regression ablation, with the largest gains on tasks where absolute scores are hardest to calibrate.

---

The chemical space relevant to drug discovery contains many more candidate molecules than can feasibly be synthesized and assayed, so the prioritization of compounds for experimental testing is a central concern of the design process [1, 2]. Machine learning has been applied to this prioritization for several decades, primarily through surrogate models that rank unseen candidates within established chemical libraries [3–7], and more recently through generative models that extend the search beyond such libraries by proposing novel candidate structures [8, 9]. The practical performance of these methods has nevertheless proven uneven, and machine-learning generators have at times been reported to underperform simple genetic-algorithm baselines on standard benchmarks [10].

While the *generation* step can draw on large, diverse datasets of chemical structures, the *scoring* of candidates against a specific downstream objective is typically constrained to the small set of measurements gathered for the target of interest, placing the scoring problem in a low-data regime [11, 12].

Scoring functions used in current generative frameworks are typically not trained on the specific biological system of interest, and therefore cannot be expected to reliably predict the activity of proposed candidates [13]. Reliable scoring in a given campaign instead requires system-specific experimental data, which is typically acquired through in-vitro assays. The readouts of such assays vary in nature, ranging from continuous measurements such as IC50 values, which can be modeled with standard regression, to discrete outputs such as hit/miss labels or potency classes, often accompanied by orthogonal molecular properties of interest. Recent evidence suggests, however, that learning to predict the relative preference between two candidate molecules can be substantially more reliable in such low-data regimes than learning to predict their absolute scores [14], because the available measurements often remain adequate to identify which of two candidates is more promising even when they are insufficient to train a robust regressor or classifier. Comparative selection over local edits has a long history in adjacent biological design problems, where directed evolution [15] and more recently Direct Preference Optimization of protein language models [16, 17] have shown that iterative small modifications under pairwise comparison can navigate large discrete design spaces effectively.

We propose treating small-molecule lead refinement as a preference-guided active learning problem, in which the surrogate at the heart of the loop is trained on relative comparisons between candidate molecules rather than on absolute predicted scores, and in which the choice of molecular generator is decoupled from the surrogate’s mechanism for scoring and ranking. We develop APOSM (Active Preference Optimization for Small-Molecule design) as a concrete instantiation of this paradigm. At each round, APOSM proposes new candidate molecules, ranks them with a learned pairwise preference surrogate, and queries an oracle on the highest-ranked subset, with the surrogate retrained on the new measurements between rounds. The framework is structured around the batched acquisition cycles typical of wet-lab screening, and the preference layer is generator-agnostic. Our implementation uses a fragment-based molecular editor [18] drawing from the ChEMBL database [19, 20], but any molecular generator could be used in its place.

We evaluate APOSM in two complementary settings. The first is a GPCR ligand rediscovery task using real-world dopamine-receptor data from GPCRdb [21, 22]: given a held-out target ligand and a warm-start library of 300 prior molecules, the optimizer must recover structurally similar high-affinity compounds. The second is the Practical Molecular Optimization (PMO) benchmark of 25 cold-start tasks spanning similarity, multi-parameter optimization, rediscovery, scaffold-hopping, docking, and individual property objectives [23, 24].

Across both settings, APOSM improves target attainment and sampling efficiency over an unguided fragment-based baseline, the Graph-GA genetic algorithm, and a pointwise-prediction ablation, with the largest gains in task groups where calibrated absolute scores are hardest to learn.

## Results

### Preference-guided refinement of small-molecule leads

APOSM operates as an active learning loop over a population of candidate molecules, repeated for a fixed number of rounds within a given oracle budget. At each round the current population is expanded by proposing local structural perturbations of its members, the resulting candidates are scored by a learned pairwise preference surrogate, the most promising subset is sent to the oracle for evaluation, and the surrogate is retrained on the augmented set of oracle-labelled molecules before the next round begins. The batched structure of the loop matches the cadence of wet-lab screening. The three components of each round, generation, scoring, and ranking, are described below, and the formal pseudocode of the full loop is given as Algorithm 1 in Methods (Sec. 3).

#### Generation: Fragment-based perturbation

Each round begins by expanding the current population of parent molecules into a larger set of candidate offspring. The framework is generator-agnostic, requiring only an operator that takes a parent population and returns a valid offspring population. For the lead-refinement setting we focus on, the operator should be local, producing offspring that remain close to their parent in chemical space, since this preserves scaffolds and core functional groups across rounds and keeps the optimization trajectory chemically plausible rather than drifting into regions of low synthetic accessibility. We instantiate the generator with the fragment-based mutation and growth procedure CReM [18], which replaces a connected sub-fragment of a parent molecule with a fragment drawn from a curated library, subject to a context-matching constraint at the attachment sites parameterized by a context radius *r*. We use *r* = 1, the most permissive setting, which admits any context-consistent substitution. The fragment library is derived from the ChEMBL database [19, 20], so the perturbations remain in the region of chemical space populated by known bioactive molecules. The formal CReM operator definition is given in Methods (Sec. 3).

#### Scoring: Pairwise preference surrogate

Generated candidates are scored against each other by a learned pairwise preference surrogate. The surrogate is a message-passing graph neural network (MPGNN) with a shared encoder that is applied to both molecular graphs in an input pair (*s*_*i*_, *s*_*j*_), followed by a preference head that consumes the pair embedding and returns a single scalar logit *z*_*ij*_. We supervise the surrogate against the sign of the oracle delta between the two molecules in the pair. With *y*(*s*) denoting the oracle score for molecule *s* and the delta defined as Δ*y*_*ij*_ = *y*(*s*_*i*_) − *y*(*s*_*j*_) the surrogate is trained on binary preference labels *t*_*ij*_ = **1**[Δ*y*_*ij*_ *>* 0] using a binary cross entropy loss on the predicted logit, ℒ := −*t*_*ij*_ log *σ*(*z*_*ij*_) − (1 − *t*_*ij*_) log(1 − *σ*(*z*_*ij*_)), which at its optimum recovers the preference log-odds

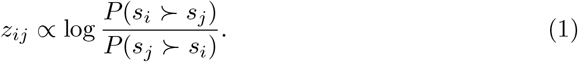

The sign of the predicted logit indicates which molecule is preferred and its magnitude reflects the surrogate’s confidence. Training pairs are sampled uniformly from the set of oracle-labelled molecules accumulated up to the current round, near-ties with |Δ*y*_*ij*_| below a small threshold are excluded, and a held-out validation set is used for early stopping. The MPGNN encoder, the pair feature, and the training procedure are specified in Methods (Sec. 3). As an ablation, we also evaluate a pointwise surrogate that predicts absolute molecule scores *ŷ*(*s*) via standard regression with Chemprop [25] and converts predictions to pairwise deltas by subtraction, 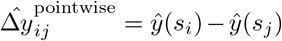. This baseline isolates the contribution of preference supervision with the rest of the loop held fixed.

#### Ranking: Global Bradley-Terry order over predicted preferences

The surrogate produces pairwise preference logits, but the acquisition step requires a single global ordering of all candidates. We obtain this ordering with a Bradley-Terry model [26]. From the predicted preference logits we construct a dominance set 𝒟_≻_ of ordered pair indices (*i, j*) for which the surrogate predicts *s*_*i*_ ≻ *s*_*j*_, and we estimate the latent ranking parameters *π* by maximizing the log-likelihood

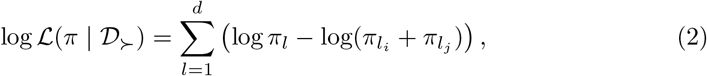

where the sum runs over the *d* observed pairwise dominance relations and *π*_*l*_ denotes the ranking parameter of the winner of comparison *l*. Because the candidate set can contain on the order of 10^5^ molecules per round, we use the Iterative Luce Spectral Ranking algorithm [27], which computes the same maximum-likelihood estimate via a stationary Markov chain construction and converges substantially faster than the classical majorization-minimization procedure [28]. To control the cost of the global fit further, we use a two-stage architecture in which each parent first generates a shortlist of its top-*K* children under per-parent surrogate predictions, and the global Bradley-Terry fit is run only on the resulting shortlist, with a diversity cap that limits any single parent’s contribution to at most 25% of the next-round population. The candidates ranked highest by the global 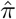 are sent to the oracle.

### Preference-guided exploration efficiently recovers GPCR ligands

We evaluate APOSM against three baselines that we apply consistently across the empirical comparisons below. These are unguided fragment-based optimization (raw CReM expansion without surrogate guidance), the Graph-GA genetic algorithm [10, 29], which has been reported as a strong baseline for molecular generation, and a pointwise-surrogate ablation that replaces the pairwise preference surrogate with a Chemprop regression model predicting absolute molecule scores.

We begin with a GPCR ligand rediscovery task using small-molecule ligands binding to a dopamine receptor [22]. The task objective is to recover a high-affinity compound given a randomly selected starting ligand, scored by Tanimoto similarity to the held-out target on extended-connectivity fingerprints, with each method given an initial training pool of 300 known dopamine-receptor ligands as warm-start supervision.

Figure 2A plots the trajectory of each method through a Tanimoto-derived two-dimensional projection of chemical space across seven generations, with the fitness landscape overlaid as the vertical coordinate. Graph-GA covers a broader region of chemical space than APOSM, but its coverage is largely peripheral to the high-fitness ridge. APOSM tracks more closely along the high-fitness region and reaches higher mean scores per generation and a higher cumulative best-so-far value (Fig. 2C). The intra-generation diversity of APOSM’s proposals is correspondingly lower than Graph-GA’s (Fig. 2B), reflecting an exploration concentrated around the active region rather than spread across the broader space. Unguided fragment-based optimization drifts away from the target ligand across generations, showing that fragment-based perturbation alone, without the preference surrogate, is insufficient to navigate toward the held-out target.

**Fig. 1:**
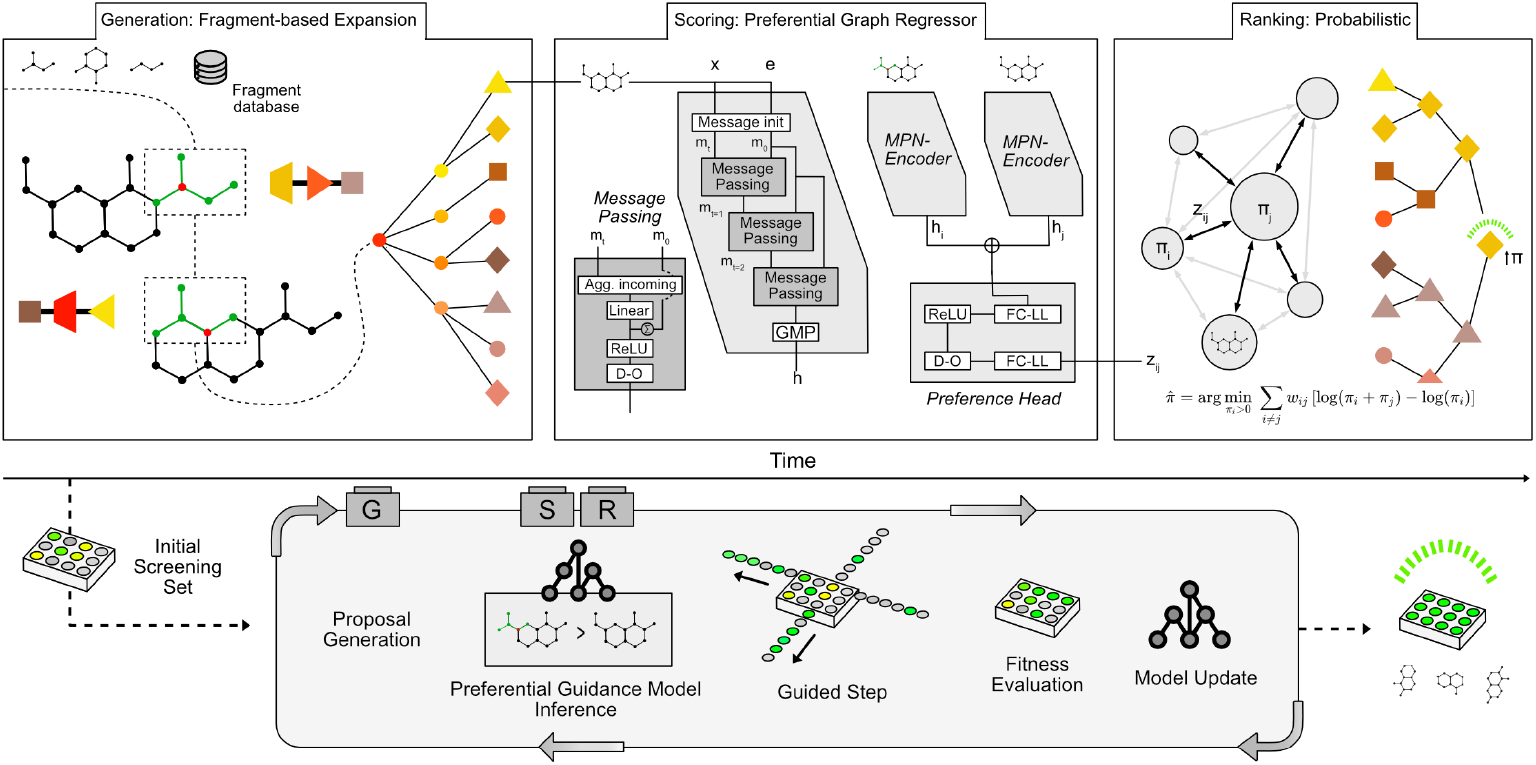
Overview of APOSM. Given an initial set 𝒟_0_ of oracle-labelled molecules and a fragment database, the active learning loop proceeds in rounds. At each round, the fragment-based generator (CReM) proposes local structural perturbations of the current candidate population, the pairwise preference surrogate scores pairs of resulting candidates, a global Bradley-Terry ranking aggregates these predicted preferences, and the oracle is queried on the top-ranked subset. The oracle-labelled molecules are added to the training set and the surrogate is retrained before the next round. The surrogate is a message-passing graph neural network with a shared encoder applied to both molecular graphs of an input pair, followed by a preference head that returns a single scalar logit.

**Fig. 2:**
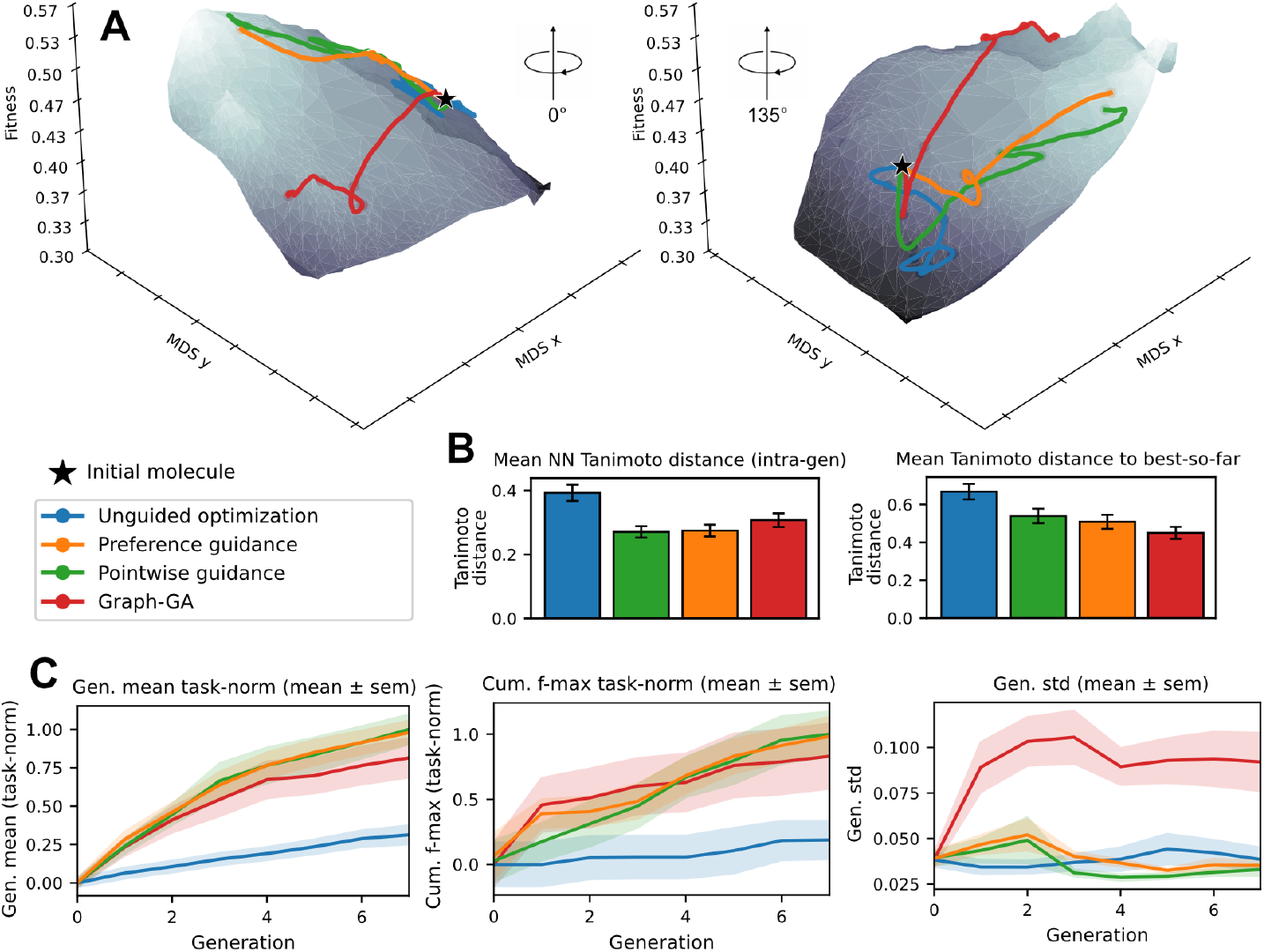
GPCR dopamine-receptor ligand rediscovery: chemical-space trajectories and performance metrics. **A**: MDS projection of generated molecules over seven generations, coloured by fitness. Trajectories show the mean position per generation. **B**: Within-generation diversity, measured as mean nearest-neighbour Tanimoto distance (left) and mean Tanimoto distance to the best-so-far molecule (right). **C**: Per-seed performance across five seeds, normalized by the per-task observed maximum. APOSM reaches higher mean and *f* -max scores than Graph-GA and unguided optimization, with a lower score standard deviation reflecting a more focused trajectory through chemical space.

The GPCR task is constrained by the size of the curated ligand dataset, with the 300-molecule pretraining set drawn from the same pool as the held-out targets. To stress-test APOSM in a fully cold-start setting across a broader range of objectives, we now turn to the Practical Molecular Optimization (PMO) benchmark.

### Preference guidance improves compound optimization on PMO

The PMO benchmark [23, 24] comprises 25 tasks spanning similarity, multi-parameter optimization, rediscovery, scaffold-hopping, docking, and individual physicochemical property objectives. Each task is initialized from a single unscored molecule and constrained to a total budget of 1000 oracle evaluations, placing the search firmly in the low-data regime relative to the combinatorial chemical space available for exploration.

Figure 3A and Supplementary Figure G2 show that both APOSM and the pointwise-surrogate ablation outperform unguided fragment-based optimization and Graph-GA on two complementary metrics, namely the mean score of generated candidates within each generation, which captures the immediate quality of proposals, and the cumulative best-so-far across the run, which captures the strongest candidate identified by the optimizer at each oracle budget level. Aggregated across task groups (Table 1 and SI Sec. G), APOSM outperforms Graph-GA on similarity, multi-parameter optimization, rediscovery, scaffold-hopping, and docking, and matches it on basic property optimization. Compared to unguided fragment-based optimization, preference guidance improves performance in every task group, which isolates the surrogate as the source of the improvement since the guided and unguided variants share the same CReM generator.

**Table 1:**
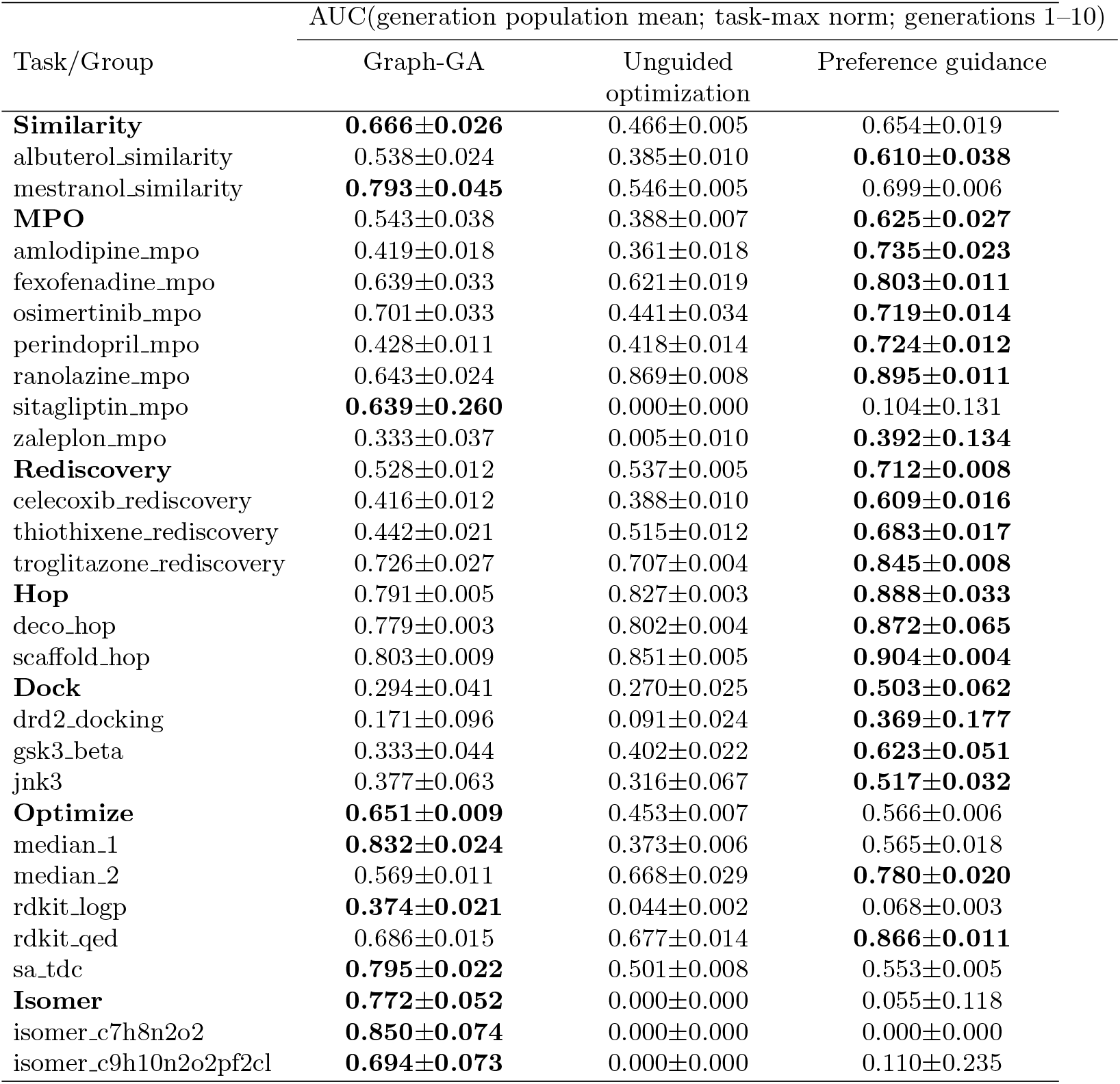
Generation-wise task-max normalized AUC of population mean per generation (*±*std across seeds). Rows include group aggregates (bold) and per-task values.

| Task/Group | AUC(generation population mean; task-max norm; generations 1–10) |  |  |
| --- | --- | --- | --- |
|  | Graph-GA | Unguided optimization | Preference guidance |
| **Similarity** | **0.666±0.026** | 0.466±0.005 | 0.654±0.019 |
| albuterol_similarity | 0.538±0.024 | 0.385±0.010 | **0.610±0.038** |
| mestranol_similarity | **0.793±0.045** | 0.546±0.005 | 0.699±0.006 |
| **MPO** | 0.543±0.038 | 0.388±0.007 | **0.625±0.027** |
| amlodipine_mpo | 0.419±0.018 | 0.361±0.018 | **0.735±0.023** |
| fexofenadine_mpo | 0.639±0.033 | 0.621±0.019 | **0.803±0.011** |
| osimertinib_mpo | 0.701±0.033 | 0.441±0.034 | **0.719±0.014** |
| perindopril_mpo | 0.428±0.011 | 0.418±0.014 | **0.724±0.012** |
| ranolazine_mpo | 0.643±0.024 | 0.869±0.008 | **0.895±0.011** |
| sitagliptin_mpo | **0.639±0.260** | 0.000±0.000 | 0.104±0.131 |
| zaleplon_mpo | 0.333±0.037 | 0.005±0.010 | **0.392±0.134** |
| **Rediscovery** | 0.528±0.012 | 0.537±0.005 | **0.712±0.008** |
| celecoxib_rediscovery | 0.416±0.012 | 0.388±0.010 | **0.609±0.016** |
| thiothixene_rediscovery | 0.442±0.021 | 0.515±0.012 | **0.683±0.017** |
| troglitazone_rediscovery | 0.726±0.027 | 0.707±0.004 | **0.845±0.008** |
| **Hop** | 0.791±0.005 | 0.827±0.003 | **0.888±0.033** |
| deco_hop | 0.779±0.003 | 0.802±0.004 | **0.872±0.065** |
| scaffold_hop | 0.803±0.009 | 0.851±0.005 | **0.904±0.004** |
| **Dock** | 0.294±0.041 | 0.270±0.025 | **0.503±0.062** |
| drd2_docking | 0.171±0.096 | 0.091±0.024 | **0.369±0.177** |
| gsk3_beta | 0.333±0.044 | 0.402±0.022 | **0.623±0.051** |
| jnk3 | 0.377±0.063 | 0.316±0.067 | **0.517±0.032** |
| **Optimize** | **0.651±0.009** | 0.453±0.007 | 0.566±0.006 |
| median_1 | **0.832±0.024** | 0.373±0.006 | 0.565±0.018 |
| median_2 | 0.569±0.011 | 0.668±0.029 | **0.780±0.020** |
| rdkit_logp | **0.374±0.021** | 0.044±0.002 | 0.068±0.003 |
| rdkit_qed | 0.686±0.015 | 0.677±0.014 | **0.866±0.011** |
| sa_tdc | **0.795±0.022** | 0.501±0.008 | 0.553±0.005 |
| **Isomer** | **0.772±0.052** | 0.000±0.000 | 0.055±0.118 |
| isomer_c7h8n2o2 | **0.850±0.074** | 0.000±0.000 | 0.000±0.000 |
| isomer_c9h10n2o2pf2cl | **0.694±0.073** | 0.000±0.000 | 0.110±0.235 |

**Fig. 3:**
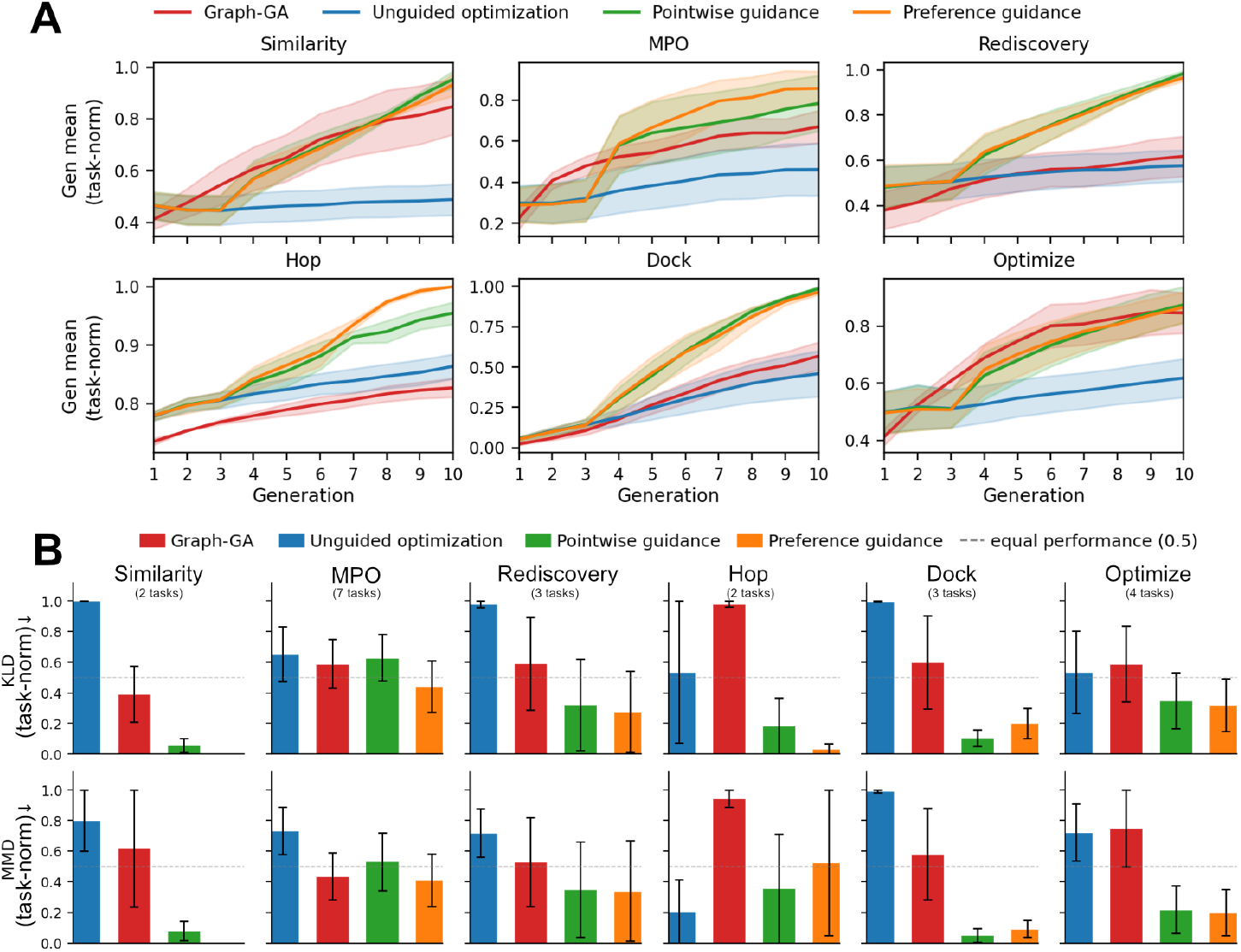
PMO benchmark performance and target-distribution alignment across task groups. **A**: Per-generation mean score, averaged within each task group (Similarity, MPO, Rediscovery, Hop, Dock, Optimize) and normalized by the per-task observed *f* - max. Shaded bands are *±*SEM across tasks. The `rdkit_logp` task is excluded; isomer and ungrouped task groups are not displayed. **B**: Per-group divergence between each method’s score distribution and the score distribution of the top 25,000 ENAMINE druglike compounds per task, measured by the forward Kullback-Leibler divergence (top) and by Maximum Mean Discrepancy with an RBF kernel (bottom). Values are rank-normalized across solvers within each task before averaging within groups, so the best method on a task receives 0 and the worst receives 1. Bars show the mean across tasks in each group with the standard error indicated.

### Preference-guided proposals align with druglike target distributions

Beyond the absolute task performance, we ask whether APOSM’s proposals also fall closer to a reference distribution of druglike synthesizable candidates. We compute divergences between the empirical score distribution of each method’s proposals and the score distribution of the top 25,000 ENAMINE druglike compounds per task (see Methods). Figure 3B shows that preference guidance reduces both the forward Kullback-Leibler divergence and the Maximum Mean Discrepancy against this reference on every task group except multi-parameter optimization (MPO), indicating that the guided proposals are closer in score distribution to the highest-scoring synthesizable candidates in the reference set than the unguided fragment-based or Graph-GA baselines. A plausible explanation for the MPO exception is that the composite nature of the objective compresses the score distribution and makes both the divergence target and the preference signal less directly informative than for the single-property task groups.

### Pairwise preference supervision outperforms pointwise regression at the surrogate level

The PMO results show that preference guidance outperforms unguided fragment-based optimization and Graph-GA, but the comparison between the pairwise preference surrogate of APOSM and the pointwise Chemprop ablation within the optimization loop is closer (Table G3). To examine the surrogate-level difference between the two formulations directly, independently of how each surrogate is used inside the loop, we performed a pre-optimization validation sweep that quantifies each surrogate’s ranking accuracy under controlled out-of-domain conditions.

For each PMO task we generated a diverse pool of molecules by mutating the seed molecule with a permissive CReM expansion, split the pool into a training set and a held-out set, and evaluated each surrogate on pairs drawn from the held-out set under the OOD-1 protocol, in which one molecule of each evaluation pair is absent from the training pairs (see Methods). This setup eliminates transitive information leakage through shared molecules and provides a stringent test of how well each surrogate generalizes to unseen comparisons. We report pairwise sign accuracy and Spearman rank correlation between predicted and true score deltas (Supplementary Fig. E1). The pairwise preference surrogate outperforms the pointwise Chemprop ablation on most PMO tasks, with comparable performance on roughly half the tasks, and the size of the advantage tracks task difficulty, with the largest gains on the tasks where absolute regression is hardest to learn.

## Discussion

The main finding of this work is that pairwise preference learning is a more reliable surrogate target than absolute regression in the low-data molecular regimes typical of screening campaigns, and that this reliability translates into improved optimization efficiency when the surrogate is placed inside an active acquisition loop. Three complementary lines of evidence support this claim. On the 25 tasks of the PMO benchmark, APOSM outperforms unguided fragment-based optimization and a strong genetic-algorithm baseline across most task groups, and matches or exceeds the pointwise-regression ablation on the integrated optimization metric. At the surrogate level, the preflight out-of-domain validation sweep shows that pairwise preference prediction is more accurate than absolute regression on the majority of PMO tasks, with the largest gains on the tasks where absolute calibration is hardest. On the GPCR ligand rediscovery task, where supervision is derived from realistic assay-style comparisons against a warm-start library of 300 dopamine-receptor ligands, APOSM tracks the high-fitness ridge of chemical space more closely than the unguided fragment-based or genetic-algorithm baselines. Across these three settings, preference-based supervision provides a more reliable surrogate signal than absolute regression for iterative optimization with noisy and sparse measurements.

The closest prior work explores pairwise prediction without active acquisition. Fralish et al. [14] and Wang and King [30] both show that pairwise architectures outperform absolute regression on small molecular datasets, but do not deploy their surrogates inside an optimization loop; thus not addressing whether the prediction-level advantage carry through to better candidate generation. Preferential Bayesian optimization [31] formalizes optimization under preference-only access but operates on fixed candidate sets without a generative component. The CheapVS framework [32] couples chemist-elicited preferences with a docking-based affinity model in an active loop but is restricted to a fixed virtual library and uses preferences as a way to capture human intuition rather than as a way to extract a more reliable signal from noisy quantitative measurements. APOSM occupies the intersection of these three threads, in that it is generative rather than fixed-library, it operates under preferences derived from quantitative measurements rather than from human elicitation, and it places the pairwise preference surrogate inside an active acquisition loop. The conceptual contribution therefore lies in how the three components fit together for the small-molecule lead refinement setting rather than in any single component on its own.

The empirical evaluation in this work concentrates on the regimes in which preference-based supervision is most naturally motivated. The PMO benchmark provides cold-start tasks where the surrogate must learn from scratch within an oracle budget, and the GPCR ligand rediscovery task provides a warm-start regime with a curated 300-molecule prior, which is closer to the operating point of practical lead optimization campaigns where some initial assay data is typically available. Three natural extensions of this scope each address a complementary direction. Substituting alternative molecular generators into the same active loop would empirically demonstrate the generator-agnosticism that the framework’s structure already implies. Extending preference supervision to genuinely multi-objective settings with explicit Pareto handling would address the multi-parameter optimization regime, where the score-distribution alignment we report in Figure 3B is least pronounced. Applying APOSM to real wet-lab assay data with batched laboratory cadence would close the loop between the methodological setting we describe and the experimental setting it was designed for.

The use of small local edits under comparative selection has long-standing precedent in protein engineering. Directed evolution [15], recognized by the 2018 Nobel Prize in Chemistry, established that iterative mutation and selection can produce highly optimized protein variants by repeatedly retaining the better-performing members of each generation. ML-guided directed evolution generalized this strategy with learned fitness surrogates [33], and Direct Preference Optimization [16] has more recently extended the paradigm to a setting where explicit pairwise preference comparisons drive both selection and generation, including for protein-language-model generators [17]. APOSM applies the same conceptual approach to small-molecule lead refinement, where the representational machinery of molecular graphs and fragment-based mutation differs from that of amino-acid sequences but the underlying loop structure of iterative comparative selection is the same. Taken together, this lineage suggests that comparative selection over local edits is an effective strategy for navigating large discrete design spaces, and the present work adds the small-molecule case to the domains where it has been demonstrated empirically.

## Methods

### Molecular distance and scoring metrics

To assess the chemical space exploration of the generative methods, we require a quantitative distance-metric between molecules. Common approaches include fingerprint-, graph-, or deep-learning derived metrics [34–36].

#### Fingerprint-based metrics

An easily-computable distance metric based on fixed-length molecular feature vectors, often binary, e.g. extended connectivity fingerprints or MACCS keys [37–39]. Chemical similarity is commonly measured as the Tanimoto-coefficient^1^ between binary representations [40], which is computable even for large compound libraries. However, from a chemist’s perspective, fingerprint bits are an abstraction of structural features, which can lead to unintuitive similarity associations [41]. In SI section G.5, we explore mean nearest neighbour Tanimoto distance as well as the tanimoto distance from the initial population, measuring the diversity within a proposed set and the novelty of the proposed set, respectively.

#### Chemical space coverage

To evaluate the chemical space covered by the generative process we use complementary metrics for property values and chemical diversity. The coverage can be computed from i) the properties of the input sets, and ii) the distributions of both labels and properties. For input set properties, distances can be computed based on fingerprint vectors (see above), which includes the mean Tanimoto similarity within a set or distance to the best observed sample or set of samples. Any *relative* comparison, e.g. distance between properties or label distributions, requires some reference set. This can be one or multiple small molecule datasets: CHEMBL [19, 20], PubChem [42], ENAMINE Real [43]; or any combination of the considered sets and their subsets. Specific applicable distribution measures are, for example, maximum mean discrepancy (MMD) [44, 45] or divergence measures like Jensen Shannon divergence [46, 47] or the Wasserstein distance [48].

#### Reference-relative label divergences

Given an ideal reference set of molecules–that is by construction or assumed to be ideal–divergences between the distribution of generated samples and the reference set can be computed. We compute divergences in the label space, e.g. by comparing the distribution of generated samples properties to those of the reference set, which should be representative of the desired chemical space. In case a generative process proposes better candidates by the target score, e.g. within the 95th percentile of observed values, we include this in the idealized reference set. We explore this approach in Section 3.

### GPCR ligand rediscovery task

To evaluate surrogate-guided optimization in a data-rich regime, we introduce a GPCR rediscovery task. At the start of each run a target molecule is randomly sampled from the reference set, then removed from the pool; a subset (n=300) of the remaining ligands form the initial starting population of scored molecules available to the optimizer. The objective is to maximize Tanimoto similarity (ECFP4, radius 2, 2048 bits) between generated candidates and the held-out target, yielding scores ∈ [0, 1]. Unlike the PMO benchmark, where all methods start from a single unscored molecule, this warm-start scenario provides the surrogate models with an immediately informative training set, directly testing whether guidance can exploit prior structure-activity data.

### Practical Molecular Optimization: In silico oracles to evaluate novel compounds

To evaluate our generative algorithm, we rely on several oracles for small molecules. An oracle, in this context, is an easy to evaluate function of any generated input. It serves as a proxy for an experiment (e.g. in-vitro) validation, which are bottlenecks when testing a conditionally generative method. Therefore, the proposed oracles serve as a *black-box* function assigning a computable score to an input molecule. The PMO benchmark is a collection of such oracles, each defining a distinct *problem*. Solving a *problem* in this space amounts to learning the mapping from an input domain to a label domain, the oracle effectively defines the entire observable problem space.

### Pairwise molecular preference surrogate

#### Pair feature and preference head

The shared message-passing encoder produces a single embedding *h*_*s*_ ∈ ℝ^*d*^ for each molecular graph *s*. For an input pair (*s*_*i*_, *s*_*j*_) the two embeddings are combined into a

pair feature that is fed to a preference head, a MLP with a single hidden layer followed by a linear projection to one output. The pair feature construction and the encoder architecture are given in detail in SI Sec. D.1; the message-passing layers are specified in Section 3.

#### Training procedure

At each round we sample unique pairs from the set of oracle-labelled molecules accumulated up to the current round. Pairs with |Δ*y*_*ij*_| below a small tie threshold *ε*_Δ_ are excluded so that the preference label is well-defined. A held-out validation subset of pairs is used for early stopping under the binary cross entropy loss with logits. Specific hyperparameters (batch size, learning rate, maximum number of training pairs, early-stopping patience) are listed in SI Table D1.

#### Pointwise Chemprop ablation

The pointwise ablation replaces the pairwise preference surrogate with a Chemprop [25] model trained to predict single-molecule oracle scores *ŷ*(*s*) ≈ *y*(*s*) under a mean-squared-error objective on continuous targets. Pairwise preferences are induced at evaluation time by subtraction of the predicted scores, 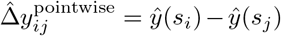. For tasks whose oracle returns a binary label, the Chemprop model is trained as a binary classifier and predicted probabilities are converted to logits before subtraction so that the ranking signal remains continuous. All other components of the active learning loop, including the generator, the acquisition step, and the ranking procedure, are identical between the pairwise and pointwise variants so that the ablation isolates the contribution of preference supervision.

#### Guided selection and ranking

When surrogate guidance is active, both the pairwise and pointwise variants employ the same two-stage ranking procedure, in which the surrogate first filters offspring locally per parent and a global Bradley-Terry fit is then performed on the shortlist. A per-parent diversity cap then limits any single parent’s contribution to at most 25% of the next-round population before the candidates are sent to the oracle. When the surrogate is not yet ready (for example during the initial warm-up rounds before enough labelled molecules have accumulated), the solver falls back to non-guided black-box selection.

#### Pre-optimization surrogate validation

To quantify the surrogate’s ranking accuracy independently of how it is used inside the optimization loop, we run a pre-optimization (*preflight*) validation sweep. For each task we generate a diverse pool of molecules with a permissive CReM expansion, split the pool into a training set and a held-out set, and evaluate each surrogate on pairs drawn from the held-out set under the OOD-1 protocol described in Sec. 3.

### Constructing train- and test-sets

We construct train and test splits as subsets derived from the set of all labeled *N* molecules, 𝒟_0_, given the training indices ℐ_train_ = *{*(*i, j*)|*i, j* ∈ [1, …, *N*]*}* such that 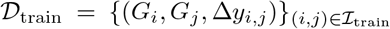, the set of molecular pairs in training. Any of the unseen inputs we denote as ℐ_test_ = ℐ_𝒢*×* 𝒢_\ℐ_train_. We evaluate the model with three tasks defined by their degree of relatedness between train and testing, i.e. out-of-distribution (OOD) scenarios. The tasks for test-set construction are

(`OOD-0`) both molecules are in the training set, but the pairs for testing are held out,

(`OOD-1`) one molecule in each test pair is not in any training pair, and

(`OOD-2`) both molecules in each test pair are not in the training set.

For details see appendix Sec. E . The `OOD-1` setting is most suitable for our use-case, the preflight surrogate validation, and the regressor is evaluated using this split.

### Probabilistic ranking over inferred scores

The Iterative Luce Spectral Ranking algorithm [27] computes the maximum-likelihood estimate of the Plackett-Luce model, which operates on general partial rankings. In the special case where each partial ranking contains exactly two items, the Plackett-Luce model coincides with the Bradley-Terry model and the log-likelihood reduces to the Bradley-Terry log-likelihood for pairwise comparisons. Compared with the classical Bradley-Terry majorization-minimization procedure [28], ILSR uses a stationary Markov chain construction to obtain a global parameter estimate, which converges substantially faster and is numerically more stable on the comparison volumes APOSM generates, on the order of 10^5^ pairs per round. A derivation of the sequential two-stage Bradley-Terry equivalence used by APOSM is given in SI Sec. F.

### Algorithm

We combine fragment-based generation with the pairwise structure-activity estimator, and the preference ranking model to iteratively refine small molecule lead compounds. To limit the number of total comparisons and speed-up required inference steps we first construct a top-K pair-set for each parent^2^, followed by importance sampling on molecule pairs to a specified (*c*_max_) constant number of comparisons. Then, within the set of importance sampled offspring, initially only compared to their own parents, a second round of inference establishes *global* preferences among shortlisted offspring. For the *pairwise* algorithm we substitute Δ*ŷ* with the predicted model logits, *z*_*ij*_ instead and fit the model with a binary cross-entropy loss on preference classes, whereas the *pointwise* model prediction target is Δ*ŷ* with the training target being an MSE loss residing on the pointwise *y*_*i*_. For the pointwise guidance ablation although the notation reads model(*x*_*i*_, *x*_*j*_), there is in fact a direct score prediction *ŷ*(*s*_*i*_), such that the function and model space may be the simpler 𝒢 *1*→ ℝ.

#### Variable annotation

𝒟_0_ initial set, *b*_0_: batch size for initial set, |*D*_*t*_|: budget for generation at time t, *b*_*f*_ : (max) batch size during optimization, 𝒟_*comp*_: Union set (over offspring) of top-k pairwise comparisons per parent, Δ*ŷ*: *global* predicted preference logits/distances (depending on surrogate type), 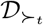: Binary preference labels for Bradley Terry/ISLR ability estimation. *ϵ*_Δ_: threshold/tolerance filter (on either predicted differences or preference logits), *k*: Population size.

##### Algorithm 1 Preference-guided molecular refinement

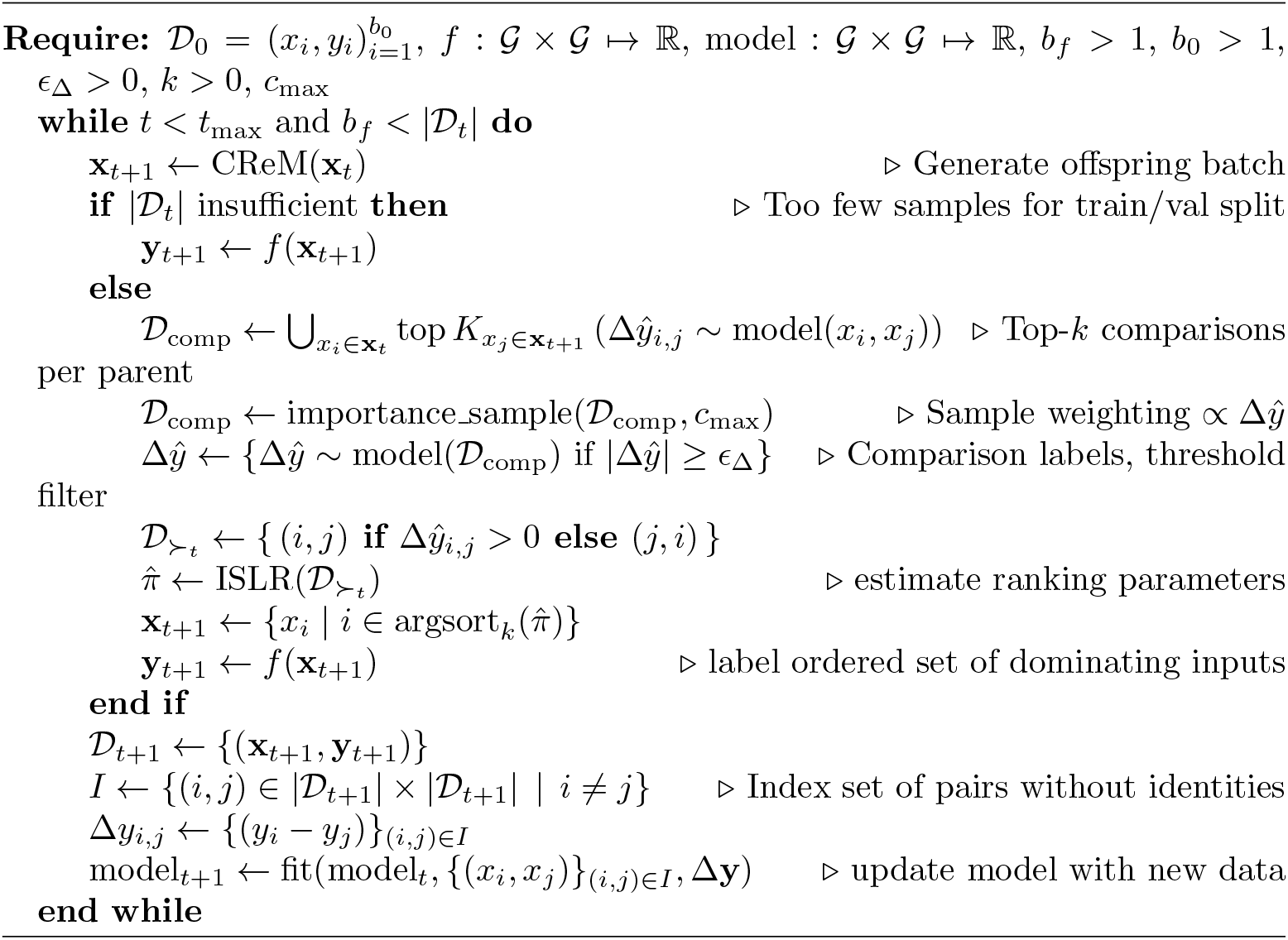

### Message passing graph neural network encoder

We represent each molecule as a graph *G* = (*V, E*), where *V* is the set of atoms and *E* is the set of bonds. Each atom *v* ∈ *V* is associated with features 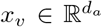, and each bond (*u, v*) ∈ *E* with features 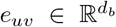. We adopt a directed, bond-centered message passing scheme, where messages are defined on directed edges. Both surrogate inputs use the same molecular graph representation, which is obtained from a SMILES string converted, via RDKit, to atom and bond features. Atom features encode element identity, valence/degree state, formal charge, chirality, hydrogen count, hybridization, aromaticity, and a mass-related channel; bond features encode bond type, conjugation, ring membership, and stereochemistry. The encoder propagates information along directed bonds and learns a fixed-size molecular embedding, from which we construct a pair feature, which is input to the *preference head*, see SI sec. D.1.

#### Message initialization

The initial edge messages in the MPGNN encoder are computed via a learned linear transformation followed by a nonlinearity:

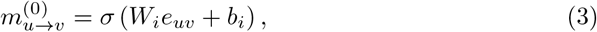

where 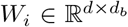 and *σ* is a nonlinear activation function (ReLU in our implementation).

#### Message passing

Messages are iteratively updated for *T* steps. At each step *t*, messages are aggregated from neighboring edges and updated as:

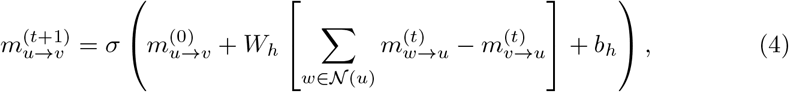

where *N* (*u*) denotes the neighbors of atom *u*. The subtraction term removes the reverse message 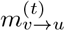, preventing immediate backtracking along edges. The residual connection to 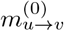 stabilizes message propagation and preserves initial edge information.

#### Atom representation

After *T* message passing steps, atom embeddings are constructed by aggregating incoming messages and combining them with atom features:

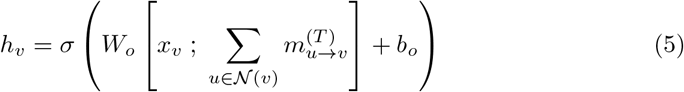

where [·;·] denotes concatenation. This combines intrinsic atom features with learned neighborhood context.

#### Readout

A fixed-dimensional molecular representation is obtained via a permutation-invariant readout over atom embeddings. We use global mean pooling:

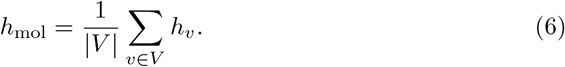

The encoder propagates information along directed bonds through iterative neighborhood aggregation and residual updates anchored to the initial messages. The resulting atom representations are pooled to produce a fixed-size molecular embedding.

### CReM – Chemically Reasonable Mutations

Given a parent set of molecules Pa = *{*(*V*_Pa_, *E*_Pa_)*}*, a reference library from which we can derive a fragment set ℱ = *{*(*V*_*F*_, *E*_*F*_)*}*, and a radius for replacement operations *r* ∈ ℕ, we can define a replacement operator which applies a fragment *F* ∈ ℱ to a parent to generate an offspring set *C* by the operation *O*_*r*_ : (Pa, *F*) −→ *C* A connected sub-fragment *F*_0_ ⊆ Pa whose removal opens *k* sites is chosen. In the special case of a grow-operation, select a heavy-atom *a* ∈ *V*_Pa_^3^ and set *F*_0_ = {H} . Select a library fragment *F* ∈ ℱ for which the context of the *k* attachment atoms matches the *k* open sites at radius *r*. Set *F* into Pa at the open sites to form the child molecule *C* ∈ *C*.

Let the intersection graph be defined as

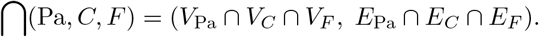

The context is legal if, for all attachment atoms *a* ∈ Pa, the neighborhood subgraph 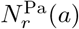, defined as the subgraph induced by all atoms within topological distance *r* from *a* in the parent, is contained in the intersection,

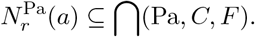

The value of *r* ∈ ℕ is the context radius, and determines the size of the chemical environment required to match between parent, child, and fragment during the operation. If this condition is satisfied, then *O*_*r*_(Pa, *F*) yields a chemically valid child. The grow-operator *F*_0_ = {H} attaches any mono-dentate^4^ fragment at an implicit hydrogen.

Basing the generative components on `CReM` naturally constrains us to chemically valid proposals with enhanced synthetic plausibility [18]. This deliberate choice reflects that the most valuable proposals, to us, are practically synthesizable candidates, which remain close to an initial molecule set. This preserves key scaffolds and functional groups [49, 50].

### Graph-GA – molecular genetic algorithm

Previous investigations have shown favorable performance of this algorithm [24]; making this our baseline. The graph-based genetic algorithm (`Graph-GA`) starts with an initial population of molecules as molecular graphs. All evolutionary operations— crossover and mutation—act directly on these graphs. Akin to other genetic algorithms, the evolution is governed by a scoring function, which assigns a score to any offspring at every generation. Parents are selected stochastically so that *fit* molecules have a higher chance of reproduction, while sampling the whole population. To breed, the algorithm cuts a single (non-ring) bond in each parent and swaps the resulting fragments. After a crossover, each child may also undergo a single random edit, i.e. adding or deleting a terminal atom, changing the atom type, inserting or removing a bond, or altering a bond order. The modification undergoes validity checks. The population is updated with the highest-fitness children, while retaining a small population of the best-scoring parents – retention set size is a hyperparameter. The cycle of sampling parents, crossover, mutation, and population update repeats for a predefined number of generations or until convergence.

### Step-wise and generation-wise assessment

The `Graph-GA` implementation [10] that we employ as a baseline retains a small set of top-scoring parents across generations and applies internal validity filtering and deduplication to each offspring batch, so the number of *new* oracle calls per generation can fall below the configured offspring target. While the optimal trade-off between *when* to query an oracle and *how many* queries to issue at any single point in time is a research question in its own right, we hold the population size, oracle budget, number of generations, and offspring count fixed across all methods. To ensure a fair comparison under these constraints, we report results both step-wise (cumulative number of oracle calls) and generation-wise (per batched iteration).

### Assessing losses for implicit generative models

The generative model and subsequent algorithm we propose do not explicitly specify a distribution and therefore no tractable likelihood – making it an *implicit* generative model (see Eq. 2 [51]). Therefore, to assess the generative distributions we rely on the losses introduced in Mohamed and Lakshminarayanan (2016). We assess the quality of the proposal samples from the generative model *G*(**z**; *θ*) by either joint density differences or ratios between the generated distribution *q*_*θ*_(**x**) and the true target *p*^∗^(**x**). Here *G* denotes the generative procedure parameterized by weights *θ* and a fragment set *c*, and *q*_*θ*_(**x**) is the resulting distribution over molecules

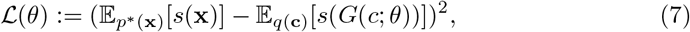

relating our assessment to moment matching of the generated sample statistics, which we determine to be the properties of the samples (Eq. 24 [51], [52]). We can formulate assessment objectives via proper scoring rules [53] or divergence measures [54]. To avoid reliance on probability densities derived from additional classifiers and the resulting modeling decisions, we assess the divergence and statistics of the generated label distributions over the samples. Specifically, we use the forward Kullback-Leibler divergence as a valid f-divergence. This leaves the question of the *assumed* true data distribution, which is challenging considering that some tasks are optimization tasks defined by the properties of the generated samples; instead of known target samples. As a reference we utilize what is readily available and synthesizable. Therefore, we use the ENAMINE REAL druglike set, and score all contained molecules with the PMO tasks, keeping the highest scoring 100 000 molecules for an individual optimization task, to obtain a valid target distribution.

## Acknowledgements

MWD and NH are supported by the NNF challenge Center for Optimized Oligo Escape and Control of Disease (NNF23OC0081287), Villum Foundation Synergy (40578), and the Swiss National Science Foundation (310030M 204518). RM is funded by the Danish Data Science Academy, which is funded by the Novo Nordisk Foundation (NNF21SA0069429) and VILLUM FONDEN (40516). WB is funded by the MLLS Center (Basic Machine Learning Research in Life Science NNF20OC0062606). Other support includes the Pioneer Centre for AI (DNRF grant number P1).

## Supplementary information

Supplementary material is available in appendix A-G

## Footnotes

1 For binary fingerprints the Tanimoto coefficient coincides with the Jaccard index; the related Dice coefficient is a distinct (though monotonically related) measure.

2 Additionally, we discard predicted preference-logits (or Δ*ŷ* for the pointwise ablation) under a small threshold as it is likely more error-prone and hence not an informative pairwise comparison.

3 Hydrogens are handled implicitly.

4 This is one attachment atom at the site.

## Notes

### Competing Interest Statement

NH is the CSO and co-founder of EDGE Biotechnologies.

