## Supplementary Material for "APOSM: Pairwise preference learning improves generative small-molecule design"

#### Appendix A Related works

Our contributions relate to (1) generative modeling for small molecules, (2) applications of *active* learning, as well as (3) preference learning for lead refinement. The objective is to propose novel molecules or refine existing candidates using machine learning (ML) algorithms [55, 56]. Hereto, Jensen introduces a graph-based *genetic algorithm* (GA) [29, 57] for molecular optimization (`Graph-GA`). Tripp and Hernández-Lobato highlights in *Genetic algorithms are strong baselines for molecule generation* [10] that minimal GAs are competitive with advanced generative ML algorithms and suggest a benchmark for molecule generators. Their findings are corroborated by the *Practical Molecular Optimization* (PMO) benchmark [24]. Here the methods REINVENT [8] and `Graph-GA`, outperform advanced algorithms with fewer evaluations. Gao et al. highlight that goal-directed molecular generation must monitor the oracle budget because in practical refinement budgets are a limiting factor.

Recent methods suggest guidance terms to traverse chemical space efficiently [58]. Applications optimize a secondary objective, e.g. synthesizability, diversity, or enhancing, for lack of a better term, the quality of a set directly with respect to a target objective [56, 59–61]. Nigam et al. present `GA-D` as molecular optimization combining a GA with a discriminator network, which steers the search away from over-exploited local maxima [62]. Zhou et al. take a *reinforcement learning* approach for generation under a graph-convolutional policy; operating directly on molecular graphs through graph-editing actions [63]. Graff and Coley employ several models in `MolPal` using molecular representations for surrogate models to explore docking as an in-silico screening tool rather than generating. Additionally, an active-learning loop refines their surrogate. Klarner et al. show that a learned data-generating stochastic process can be regularized to be guided within a context − applicable to small molecules.

Preference-learning is, at least for now, less common in the literature. Choung et al. frame optimization as a preference problem for a latent-ranking model on pairwise comparisons assessed by 35 Novartis experts [65]. Dang et al. extend preference learning into a Bayesian optimization framework together with a structure-based affinity predictor (`CheapVS`) [32]. Their method actively selects which molecules to dock and which pairs to show the chemist. Crucially, they operate on a fixed library and are not generating de novo.

#### Appendix B Additional measures in chemical space

##### Graph-based structural metrics

directly compare molecular structures as labeled graphs, which capture explicit atom connectivity. Common methods that highlight shared substructures between molecules include graph edit distance[5] and maximum common subgraph (MCS) measures [66].

[5] The number of edits required to transform one molecular graph into another.

Although these methods largely align with chemists' structural intuitions, they can be costly to compute.

**Deep learning-derived metrics**

measure molecules embedded in some latent space usually through deep-learning model encodings, for instance variational autoencoders (VAEs) [67, 68], graph neural networks (GNNs) [69], or large language models [70]. Metrics like manifold distances or cosine-similarities can be used to characterize the latent space and its learned chemical relationships [71]. The learned embedding can incorporate multiple molecular properties  e.g. structural, physicochemical, or biological. This allows any latent exploration to implicitly inform us about the learned chemical properties. However, the interpretation of distances can lack explainability due to its complexity and any conclusions drawn from latents are model-dependent.

**Additions to coverage**

Multiple properties can be captured by measures for a hyper-volume, which covers all properties relative to a reference set, e.g. a starting- or target-set [72, 73]. Additionally, the *novelty* quantity is an established metric which measures how unique proposals are relative to some known molecule candidate set.

Proposal sets can be characterized by structural variations quantifiable by occurrences of scaffolds, rings, and functional groups (BRICS). Specifically, Murcko-scaffolds or maximum common subgraphs (MCS) allow us to quantify the diversity of a molecule set.

### Appendix C Preference logits in a BCE context

Given oracle score $y(s)$ for molecule $s$, the oracle delta for pair $(s_i, s_j)$ is

$$\Delta y_{ij} = y(s_i) - y(s_j), \tag{C1}$$

with the preference logit for pair $(s_i, s_j)$ being

$$z_{ij} \propto \log \frac{P(s_i \succ s_j)}{P(s_j \succ s_i)} \propto \log \frac{P(\Delta y_{ij} > 0)}{P(\Delta y_{ij} < 0)}, \tag{C2}$$

which follows directly from optimizing a binary cross-entropy loss on pairwise preferences. $t_{ij} \in \{0, 1\}$ denotes the preference label indicating whether $s_i \succ s_j$, and the model predicts logit $z_{ij}$ with preference probability

$$P_\theta(s_i \succ s_j) = \sigma(z_{ij}) = \frac{1}{1 + e^{-z_{ij}}}. \tag{C3}$$

The expected binary cross-entropy objective is

$$\mathcal{L}(z_{ij}) = -p_{ij} \log \sigma(z_{ij}) - (1 - p_{ij}) \log(1 - \sigma(z_{ij})), \tag{C4}$$

where

$$p_{ij} = P(s_i \succ s_j). \tag{C5}$$

Differentiating with respect to $z_{ij}$ gives

$$\frac{\partial \mathcal{L}}{\partial z_{ij}} = \sigma(z_{ij}) - p_{ij}. \tag{C6}$$

Which at the optimum is,

$$\sigma(z_{ij}^\star) = p_{ij}. \tag{C7}$$

Applying the inverse sigmoid transformation yields

$$z_{ij}^\star = \log \frac{p_{ij}}{1 - p_{ij}} = \log \frac{P(s_i \succ s_j)}{P(s_j \succ s_i)}. \tag{C8}$$

Therefore, training with binary cross-entropy on pairwise preferences learns logits proportional to the log-odds of one molecule being preferred over another.

### Appendix D Surrogate evaluation and model specifications

#### D.1 MPN encoder and preference head

Given a pair $(G_i, G_j)$, the MPN encoder produces embeddings $h_i, h_j \in \mathbb{R}^d$. We construct a pair feature

$$\hat{h}_{ij} = [h_i,\, h_j,\, (h_i - h_j),\, |h_i - h_j|,\, h_i \odot h_j] \in \mathbb{R}^{5d},$$

and pass $\hat{h}_{ij}$ through a preference head, an MLP with a single hidden layer (Linear→ReLU→Dropout) followed by a final linear layer to one output. The scalar output is the preference logit presented to the binary cross entropy criterion.

#### D.2 Model and trainer specification

In the following, we elaborate on the optimization protocol, surrogate formulations, training criteria, and preflight evaluation procedure used for guided molecular refinement.

##### D.2.1 Study design and optimization envelope

We evaluate optimization on 25 PMO objectives, spanning physicochemical property tasks, docking-guided objectives, rediscovery and similarity tasks, multi-parameter optimization tasks, scaffold-hopping settings, and isomer-style constraints. For each task and seed, we run four solvers:

1. Graph-GA as a graph genetic algorithm benchmark,
2. Unguided mutative (via CReM) optimization as a fragment-based evolutionary baseline,

3. Preference-guided mutative optimization with surrogate-guided candidate ranking,
4. Pointwise-guided (absolute property predicted) mutative optimization as an ablation to the preference-based guidance.

All four solvers are launched under matched run-level controls,

$$G = 10, \quad C = 100, \quad P = 200, \quad B = 1000, \tag{D9}$$

where $G$ is generations, $C$ is offspring target per generation, $P$ is population size, and $B$ is the total oracle budget. This preserves budget parity at the configuration level. Each task is run over five random seeds (0–4); all reported means and standard deviations are computed across these seeds. The training set for the guidance surrogate is sampled uniformly across generations from a *ledger* kept of all oracle-inquired molecules.

**Table D1**: Hyperparameters used across all benchmark experiments (PMO and GPCR rediscovery).

| Parameter | Graph-GA | Unguided optimization | Pointwise guidance | Preference guidance |
|---|---|---|---|---|
| *Shared solver parameters* | | | | |
| Evaluation budget | 1 000 | 1 000 | 1 000 | 1 000 |
| Generations | 10 | 10 | 10 | 10 |
| Population size | 200 | 200 | 200 | 200 |
| Offspring size | 100 | 100 | 100 | 100 |
| Parent sampler | — | uniform quantile | uniform quantile | uniform quantile |
| *Mutation / offspring generation* | | | | |
| Mutation operator | crossover + mutation | CReM | CReM | CReM |
| Fragment database | — | ChEMBL 22 | ChEMBL 22 | ChEMBL 22 |
| CReM context radius | — | 1 | 1 | 1 |
| *Surrogate model* | | | | |
| Prediction target | — | — | $\hat{y}(s_i) - \hat{y}(s_j)$ | $\text{logit}(\Delta y_{ij}) \propto \log \frac{P(s_i \succ s_j)}{P(s_j \succ s_i)}$ |
| Training target | — | — | $y_i$ (pointwise) | $\mathbf{1}[y_i > y_j]$ (pref.) |
| Loss function | — | — | MSE (regression) | BCEWithLogits |
| Training epochs | — | — | 150 | 25 |
| Early stopping patience | — | — | — | 5 |
| Max training molecules | — | — | 1 200 | 600 |
| Max training pairs | — | — | — | 6 000 |
| Learning rate | — | — | Chemprop default | 0.001 |
| Batch size | — | — | Chemprop default | 256 |
| Tie epsilon | — | — | — | $10^{-8}$ |
| Min surrogate molecules | — | — | 300 | 300 |
| Diversity cap fraction | — | — | 0.25 | 0.25 |

#### Nominal vs realized offspring in MolGA

Although nominal offspring controls are matched across all solvers ($C^{\text{MolGA}} = C^{\text{CReM}} = C$), the `Graph-GA` implementation retains a small set of top-scoring parents across generations and applies internal validity filtering and deduplication to each offspring batch, so the effective number of new unique offspring can fall below the configured target in some generations ($C_g^{\text{eff}} \leq C$). Parity is therefore enforced at the level of configured population, offspring target, and total oracle budget, while the realized candidate count per generation may differ in practice. Therefore, in our evaluation, we report results both step-wise (cumulative oracle calls) and generation-wise (per iteration) to transparently reflect this dynamic.

##### D.2.2 Mutative equivalence of guided and non-guided fragment-based optimization

The non-guided and guided mutative optimization share the same chemistry engine and candidate-construction primitives: identical CReM grow and mutate operators, identical fragment context radius (radius = 1), identical sanitization and canonicalization pipeline, and identical per-parent expansion cap tied to generation offspring budget.

The proposed guided procedure differs in post-generation filtering, ranking, and shortlist throughput, and not in mutation chemistry. This distinction is important for ablation interpretation: the observed difference is an attribute of guidance quality and selection policy rather than a different mutational operator family.

##### D.2.3 Pairwise objective definitions

Let $y(s)$ denote the oracle value for molecule $s$. For a pair $(s_i, s_j)$, the true delta is

$$\Delta y_{ij} = y(s_i) - y(s_j). \tag{D10}$$

The induced pairwise preference sign is

$$\mathrm{pref}(i, j) = \mathrm{sign}(\Delta y_{ij}). \tag{D11}$$

Near ties are removed when

$$|\Delta y_{ij}| \leq \varepsilon_\Delta, \tag{D12}$$

with $\varepsilon$ denoting a task-level tie threshold.

##### D.2.4 Preference surrogate trainer

The pairwise preference surrogate directly consumes two molecules and predicts a pairwise preference logit,

$$z_{ij} = f_\theta(s_i, s_j), \tag{D13}$$

with probability

$$\hat{p}_{ij} = \sigma(z_{ij}) = \frac{1}{1 + e^{-z_{ij}}}. \tag{D14}$$

Pair labels are

$$t_{ij} = \mathbf{1}[\Delta y_{ij} > 0]. \tag{D15}$$

The pairwise training loss is BCE-with-logits,

$$\mathcal{L}_{\mathrm{pair}} = -\frac{1}{N} \sum_{(i,j)} \Big[ t_{ij} \log \hat{p}_{ij} + (1 - t_{ij}) \log(1 - \hat{p}_{ij}) \Big], \tag{D16}$$

and we interpret the pair preference score as

$$\hat{\Delta y}_{ij}^{\text{preference}} = z_{ij}. \tag{D17}$$

In the implementation, unique pairs are sampled from the current assay pool, near ties are removed, pair count is capped, and early stopping is applied based on validation BCE-with-logits.

##### D.2.5 Pointwise surrogate trainer

The pointwise surrogate model serves as an ablation to the prefential guidance and is trained directly to predict pointwise labels,

$$\hat{y}(s) \approx y(s), \tag{D18}$$

and induces pairwise deltas by subtraction,

$$\hat{\Delta y}_{ij}^{\text{pointwise}} = \hat{y}(s_i) - \hat{y}(s_j). \tag{D19}$$

**Binary-task handling**

If labels are effectively binary, training switches to classification mode. Predicted probabilities $p(s)$ are mapped to logits,

$$\ell(s) = \log \frac{p(s)}{1 - p(s)}, \tag{D20}$$

and pairwise deltas are formed as

$$\hat{\Delta y}_{ij}^{\text{pointwise}} = \ell(s_i) - \ell(s_j). \tag{D21}$$

Preserving signed ranking behavior while avoiding regression-mode failures on binary labels.

**Optimization criterion**

Training is pointwise: Regression objective for continuous targets, and binary cross-entropy objective for binary targets.

##### D.2.6 Guided ranking and candidate acquisition

When guidance is active, ranking is obtained in two stages: First, a parent-local child filtering using surrogate deltas, then a global pairwise ranking among shortlisted children with anchor comparisons.

Global ranking is solved with iterative Luce spectral ranking. A per-parent diversity cap (a single parent can not be responsible for more than 25% of the offspring population) is then enforced before oracle querying. If warm-up/readiness criteria fail, our preference-based guidance procedure falls back to non-guided black-box selection.

#### D.3 Ablation II: Preflight surrogate validation

##### D.3.1 Pairwise performance metrics

Let $\mathcal{P}$ be the set of evaluated non-tie pairs. Pairwise sign accuracy is

$$\text{PairAcc} = \frac{1}{|\mathcal{P}|} \sum_{(i,j)\in\mathcal{P}} \mathbf{1}\left[\text{sign}(\Delta y_{ij}) = \text{sign}(\hat{\Delta} y_{ij})\right]. \tag{D22}$$

Spearman correlation is evaluated between

$$\{\Delta y_{ij}\}_{(i,j)\in\mathcal{P}} \quad \text{and} \quad \{\hat{\Delta} y_{ij}\}_{(i,j)\in\mathcal{P}}. \tag{D23}$$

##### D.3.2 Preflight OOD protocol

Preflight is used to assess ranking reliability before or alongside full optimization. For each task-seed unit, we initialize from task-provided molecules, expand the molecule pool with iterative CReM-based mutations, canonicalize and deduplicate, oracle-score the generated molecules, split into train and unseen pools, and sample evaluation pairs under an explicit OOD mode, ensuring that there is no link between hold-out sets via transitivity in pairs of molecules.

Current preflight pool-generation controls are:

$$n_{\text{mol}} = 5000, \quad \text{context radius} = 1, \quad n_{\text{repl}} = 30, \quad n_{\text{parents}} = 64, \quad n_{\text{rounds}}^{\text{max}} = 300. \tag{D24}$$

Current split and evaluation controls are:

$$n_{\text{train}} = 1000, \quad n_{\text{pairs}}^{\text{max}} = 30000. \tag{D25}$$

**Criteria**

are identical to the configuration of the PMO-tasks: The preference surrogate model is trained to predict preference logits with BCEWithLogitsLoss on binary pair labels indicating whether $y_i > y_j$. Pointwise *Chemprop* model: Trained as regression on oracle scores (MSE objective), with pairwise deltas formed at evaluation by subtracting predicted (direct) single-molecule scores.

### Appendix E Train- and Test-set construction for isolated surrogate evaluation

Let all labelled $N$ ligands be $\mathcal{D}_0$ with training indices $\mathcal{I}_{\text{train}} = \{(i,j)|i,j \in [1,...,N]\}$ such that $\mathcal{D}_{\text{train}} = \{(G_i, G_j, \Delta y_{i,j})\}_{(i,j)\in\mathcal{I}_{\text{train}}}$. Let

$$\mathcal{M}_{\text{train}} = \{\, m : (m,\cdot) \in \mathcal{I}_{\text{train}} \ \vee \ (\cdot, m) \in \mathcal{I}_{\text{train}} \,\}$$

be the set of molecules appearing in at least one training pair. We define the *individual out-of-domain* test index sets as

$$\mathcal{I}_{\text{test-OOD-0}} \triangleq \{(i,j) \mid (i,j) \notin \mathcal{I}_{\text{train}},\ i \in \mathcal{M}_{\text{train}},\ j \in \mathcal{M}_{\text{train}}\}, \tag{E26}$$

$$\mathcal{I}_{\text{test-OOD-1}} \triangleq \{(i,j) \mid (i,j) \notin \mathcal{I}_{\text{train}},\ (i \in \mathcal{M}_{\text{train}} \land j \notin \mathcal{M}_{\text{train}}) \lor (i \notin \mathcal{M}_{\text{train}} \land j \in \mathcal{M}_{\text{train}})\}, \tag{E27}$$

$$\mathcal{I}_{\text{test-OOD-2}} \triangleq \{(i,j) \mid i \notin \mathcal{M}_{\text{train}} \land j \notin \mathcal{M}_{\text{train}}\}. \tag{E28}$$

By construction the three sets are disjoint: `OOD-0` contains pairs of two training-seen molecules whose pairing was held out, `OOD-1` contains pairs with exactly one novel molecule, and `OOD-2` contains pairs of two novel molecules.

#### E.1 Isolated surrogate evaluation

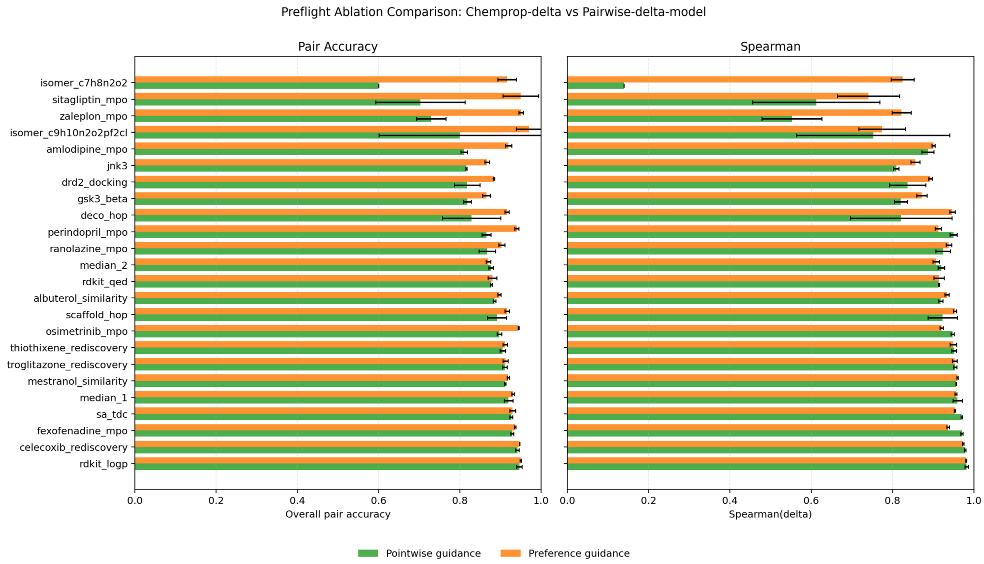


**Supplementary Fig. E1**: Comparison of estimator validation between the pairwise preference surrogate and the *Chemprop* pointwise surrogate. In the seemingly easier tasks, both surrogates perform well, while in the more difficult tasks the preference surrogate shows superior performance. The performance metrics are pairwise sign accuracy and Spearman correlation, evaluated on a held-out set of molecule pairs sampled according to the OOD-1 criterion. Spearman correlation is computed between true pairwise score differences ($\Delta y = y_i - y_j$) and the model's predicted pair score (logit for pairwise-BCE models; predicted delta for regression models). Each point represents a task, colored by task group, with the x-axis representing the pointwise *Chemprop* surrogate performance and the y-axis representing the preference surrogate performance.

### Appendix F Pairwise ranking comparisons

Instead of the proposed ILSR method for iterative ML estimate, we can also compute the Bradley Terry pairwise comparison likelihood. In practice this becomes feasible if there are sufficiently few candidates to rank. For each generation step $t$ we compute

$$\mathcal{L}_{\mathrm{BT}}^{(g)}(\beta) = \sum_{(i,j)\in P^{(g)}} \Big[ w_{i,j}^{(g)} \log \frac{e^{\beta_i}}{e^{\beta_i} + e^{\beta_j}} + \big(1 - w_{i,j}^{(g)}\big) \log \frac{e^{\beta_j}}{e^{\beta_i} + e^{\beta_j}} \Big]. \tag{F29}$$

We obtain the optimal ranking indices $\pi_t$. Maximizing $\mathcal{L}_{\mathrm{BT}}^{(g)}$ gives latent scores $\hat{\beta}^{(g)} = (\hat{\beta}_1^{(g)}, \dots, \hat{\beta}_{|\mathcal{C}^{(g)}|}^{(g)})$, where $\mathcal{C}^{(g)}$ is our offspring (children) set. The ranking permutation

$$\pi^{(g)} \;=\; \underset{i\in\mathcal{C}^{(g)}}{\operatorname{argsort}}\big[-\hat{\beta}_i^{(g)}\big] \tag{F30}$$

orders candidates from most to least promising.

#### F.1 Equivalence of a sequential Bradley-Terry model

We rank candidates in two stages: first each offspring is compared to its parent using the surrogate, then the shortlisted candidates are ranked globally (Bradley-Terry (BT) fit estimate via ILSR) . We show that the first stage requires only a sort, with no iterative estimation, and that the two-stage procedure recovers the same ranking as a single global BT fit.

##### Single-parent ranking

Consider parent $p$ with offspring $\mathcal{C} = \{c_1, \dots, c_K\}$. The surrogate is trained to approximate property differences (or property preference logits), so

$$s_{c_i} \triangleq f(G_{c_i}, G_p) \approx y_{c_i} - y_p, \quad i = 1, \dots, K. \tag{F31}$$

Because $y_p$ is the same constant for every child of the same parent, sorting by $s_{c_i}$ is equivalent to sorting by $y_{c_i}$. This requires $K$ surrogate evaluations and no iterative procedure. Moreover, the child–child comparisons are redundant: $f(G_{c_i}, G_{c_j}) \approx (y_{c_i} - y_p) - (y_{c_j} - y_p) = s_{c_i} - s_{c_j}$, so they carry no information beyond what the parent-anchored scores already provide.

##### Multiple parents and shortlisting

A generation contains $M$ parents $\{p_1, \dots, p_M\}$, each producing offspring $\mathcal{C}_m$. The two-stage procedure is:

1. **Parent-local sort.** For each parent $p_m$, compute $s_{c_{m,k}} = f(G_{c_{m,k}}, G_{p_m})$ for all $k$ and sort. Retain the top $n_m$ per parent to form a shortlist $\mathcal{S} = \bigcup_m \mathcal{S}_m$.
2. **Global BT ranking.** Compute all-pairs comparisons within $\mathcal{S}$ and fit a BT model (e.g. via ILSR [27]) to obtain global abilities $\{\hat{\beta}_i\}_{i\in\mathcal{S}}$.

Stage 1 is exact within each family (the constant-shift argument above), but scores from different families are *not* directly comparable: $s_{c_{m,k}} \approx y_{c_{m,k}} - y_{p_m}$, so cross-family ranking would rank by *improvement over parent* rather than absolute quality. Stage 2 resolves this by introducing cross-family comparisons that calibrate the scores onto a common scale.

The shortlist is safe, in the sense that the two-stage ranking agrees with a single global BT fit over all $\sum_m K_m$ candidates, whenever $\mathcal{S}$ contains every candidate that would appear in the global top-$k$. Because within-family ranking is exact, any globally top-$k$ candidate must also rank in the top of its own family, so retaining $n_m \geq k$ candidates per parent suffices.

##### Complexity

The two-stage approach requires $\sum_m K_m + \binom{|\mathcal{S}|}{2}$ surrogate evaluations with $|\mathcal{S}| \ll \sum_m K_m$, compared to $O\big((\sum_m K_m)^2\big)$ for a single global fit.

### Appendix G PMO Benchmark

We emphasize that RDKit derived direct optimization tasks like `logP`, `SA` TDC are easily exploitable and not necessarily sensible in a practical lead optimization context, as emphasized by [24]. The following are the results for all tasks, in the previous generation-wise and commonly reported step-wise generation.

- Similarity: albuterol_similarity, mestranol_similarity
- Multi-param. Optimization: amlodipine_mpo, fexofenadine_mpo, osimertinib_mpo, perindopril_mpo, ranolazine_mpo, sitagliptin_mpo, zaleplon_mpo
- Rediscover: celecoxib_rediscovery, thiothixene_rediscovery, troglitazone_rediscovery
- Change Dec./Scaff: deco_hop, scaffold_hop
- Dock: drd2_docking, gsk3, jnk3
- isomer: isomer_c7h8n2o2, isomer_c9h10n2o2pf2cl
- Optimization: median_1, median_2, (rdkit_logp), rdkit_qed, sa_tdc

#### G.1 Continuous evaluation metrics for molecular optimization

##### Stepwise vs. generational assessment

We differentiate between a *stepwise assessment* and a *generational assessment* in our optimization tasks, the latter corresponding to each iteration of a batched population. We perform continuous model evaluation on the PMO tasks during optimization, i.e., we track metrics for every batch and for every additional compound generated. This distinction is necessary because conditionally generative models for small molecules generally lack an intrinsic notion of batch size, whereas in our setting -both for active learning and the wet-lab experiments that motivated it - there is a natural batch size (e.g., dictated by synthesis or screening instrumentation).

##### Notation and indices

Let $y$ denote the observed label (score) of a generated compound. For **stepwise** assessment, we index compounds by the global step $s$ and write $y[0:s]$ for the labels of all

| Metric | Stepwise (cumulative up to $s$) | Generational (within batch $i$) |
|---|---|---|
| **Fmax** | $F_{\max}(s) = \max(Y_s)$ | $F_{\max}(i) = \max(Y_i)$ |
| **Cumul. Top 200** | $\mathrm{Top}_{200}(s) = \mathcal{T}_{200}(Y_s)$ | $\mathrm{Top}_{200}(i) = \mathcal{T}_{200}(Y_i)$ |
| **Mean** | $\mu_y(s) = \frac{1}{\lvert Y_s \rvert} \sum Y_s$ | $\mu_y(i) = \frac{1}{\lvert Y_i \rvert} \sum Y_i$ |
| **Sigma ($\sigma$)** | $\sigma_y(s) = \sqrt{\mu_{y^2}(s) - \mu_y(s)^2}$, where $\mu_{y^2}(s) = \frac{1}{\lvert Y_s \rvert} \sum Y_s^2$ | $\sigma_y(i) = \sqrt{\mu_{y^2}(i) - \mu_y(i)^2}$, where $\mu_{y^2}(i) = \frac{1}{\lvert Y_i \rvert} \sum Y_i^2$ |
| **Median** | $\mathrm{median}(s) = \mathrm{median}(Y_s)$ | $\mathrm{median}(i) = \mathrm{median}(Y_i)$ |
| **SP** | $SP(s) = \frac{\sigma_y(s)}{\mu_y(s)}$ | $SP(i) = \frac{\sigma_y(i)}{\mu_y(i)}$ |

**Table G2**: Stepwise (cumulative) and generational (per-batch) evaluation metrics used in molecular optimization.

compounds generated up to step $s$. For **generational** assessment, we index batches by $i \in \{1, 2, \ldots\}$ and let $s_i$ be the number of molecules (steps) in batch $i$; we write $y[0\!:\!s_i]$ for the labels of the compounds generated in batch $i$. We denote the mean of $y$ over a specified set by $\mu_y(\cdot)$ and the mean of squared labels by $\mu_{y^2}(\cdot)$.

##### Evaluation metrics

We report both stepwise and generational versions of the following metrics. For compactness we write $Y_s = y[0 : s]$ for the cumulative label vector up to step $s$, and $Y_i = y[0\!:\!s_i]$ for the labels within batch $i$. We let $\mathcal{T}_k(X)$ denote the mean of the top-$k$ elements of $X$ (or the mean of all elements when $|X| < k$).

Stepwise metrics provide a running view of optimization progress over time and allow direct comparison of methods with different batching or sampling schedules. Generational metrics summarize the distribution of outcomes within each batch (iteration) and align with practical constraints in active learning loops and wet-lab workflows where decisions are made at batch granularity.

All metrics are computed on the labels available at the time of evaluation. For SP and CoV, values are undefined when $\mu_y(\cdot) = 0$; when encountered, we report the other metrics and omit SP/CoV for that index. Unless otherwise stated, $\sigma_y(\cdot)$ denotes the population standard deviation computed via $\sqrt{\mu_{y^2} - \mu_y^2}$.

**Table G3**: Generation-wise task-max normalized AUC of population mean per generation (±std across seeds). Rows include group aggregates (bold) and per-task values.

| Task/Group | AUC(generation population mean; task-max norm; generations 1–10) | | | |
|---|---|---|---|---|
| | Graph-GA | Unguided optimization | Pointwise guidance | Preference guidance |
| **Similarity** | **0.666±0.026** | 0.466±0.005 | 0.662±0.025 | 0.654±0.019 |
| albuterol_similarity | 0.538±0.024 | 0.385±0.010 | 0.607±0.048 | **0.610±0.038** |
| mestranol_similarity | **0.793±0.045** | 0.546±0.005 | 0.717±0.012 | 0.699±0.006 |
| **MPO** | 0.543±0.038 | 0.388±0.007 | 0.577±0.009 | **0.625±0.027** |
| amlodipine_mpo | 0.419±0.018 | 0.361±0.018 | **0.744±0.008** | 0.735±0.023 |
| fexofenadine_mpo | 0.639±0.033 | 0.621±0.019 | **0.811±0.022** | 0.803±0.011 |
| osimertinib_mpo | 0.701±0.033 | 0.441±0.034 | 0.708±0.016 | **0.719±0.014** |
| perindopril_mpo | 0.428±0.011 | 0.418±0.014 | **0.763±0.015** | 0.724±0.012 |
| ranolazine_mpo | 0.643±0.024 | 0.869±0.008 | **0.897±0.010** | 0.895±0.011 |
| sitagliptin_mpo | **0.639±0.260** | 0.000±0.000 | 0.002±0.003 | 0.104±0.131 |
| zaleplon_mpo | 0.333±0.037 | 0.005±0.010 | 0.114±0.056 | **0.392±0.134** |
| **Rediscovery** | 0.528±0.012 | 0.537±0.005 | **0.715±0.013** | 0.712±0.008 |
| celecoxib_rediscovery | 0.416±0.012 | 0.388±0.010 | 0.601±0.028 | **0.609±0.016** |
| thiothixene_rediscovery | 0.442±0.021 | 0.515±0.012 | **0.691±0.025** | 0.683±0.017 |
| troglitazone_rediscovery | 0.726±0.027 | 0.707±0.004 | **0.853±0.005** | 0.845±0.008 |
| **Hop** | 0.791±0.005 | 0.827±0.003 | 0.870±0.031 | **0.888±0.033** |
| deco_hop | 0.779±0.003 | 0.802±0.004 | 0.846±0.061 | **0.872±0.065** |
| scaffold_hop | 0.803±0.009 | 0.851±0.005 | 0.893±0.015 | **0.904±0.004** |
| **Dock** | 0.294±0.041 | 0.270±0.025 | **0.511±0.039** | 0.503±0.062 |
| drd2_docking | 0.171±0.096 | 0.091±0.024 | **0.411±0.091** | 0.369±0.177 |
| gsk3_beta | 0.333±0.044 | 0.402±0.022 | 0.583±0.042 | **0.623±0.051** |
| jnk3 | 0.377±0.063 | 0.316±0.067 | **0.541±0.059** | 0.517±0.032 |
| **Optimize** | **0.651±0.009** | 0.453±0.007 | 0.563±0.009 | 0.566±0.006 |
| median_1 | **0.832±0.024** | 0.373±0.006 | 0.544±0.015 | 0.565±0.018 |
| median_2 | 0.569±0.011 | 0.668±0.029 | **0.787±0.040** | 0.780±0.020 |
| rdkit_logp | **0.374±0.021** | 0.044±0.002 | 0.067±0.002 | 0.068±0.003 |
| rdkit_qed | 0.686±0.015 | 0.677±0.014 | 0.866±0.012 | **0.866±0.011** |
| sa_tdc | **0.795±0.022** | 0.501±0.008 | 0.549±0.005 | 0.553±0.005 |
| **Isomer** | **0.772±0.052** | 0.000±0.000 | 0.024±0.053 | 0.055±0.118 |
| isomer_c7h8n2o2 | **0.850±0.074** | 0.000±0.000 | 0.000±0.000 | 0.000±0.000 |
| isomer_c9h10n2o2pf2cl | **0.694±0.073** | 0.000±0.000 | 0.048±0.106 | 0.110±0.235 |

#### G.2 Aggregated performance: Additional metrics

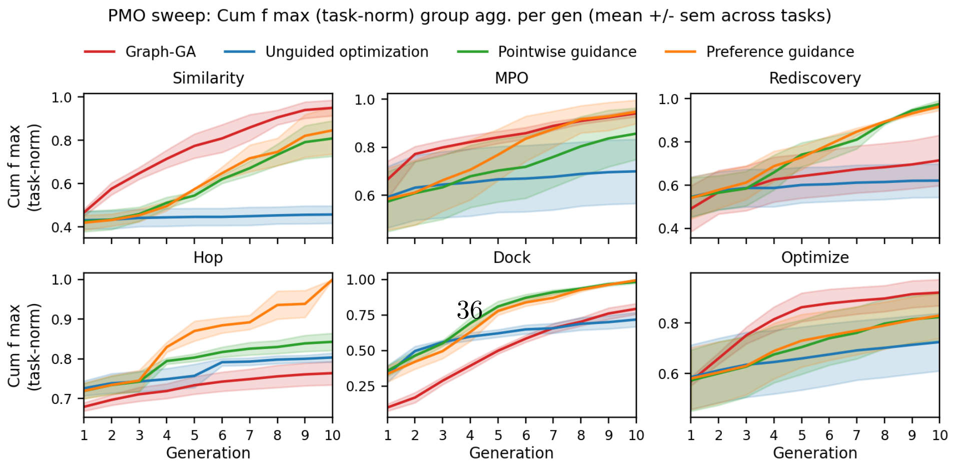


**Supplementary Fig. G2**: PMO sweep grouped aggregates for Cumulative f max (task-norm). Panels show task-group means across generations for Similarity, MPO, Rediscovery, Hop, Dock, Optimize; shaded bands are +/- sem across tasks. Excluded task(s): rdkit_logp. Groups not displayed: isomer,ungrouped. Values are normalized per task by that task's maximum observed value across all methods and generations before group aggregation.

**Table G4**: Stepwise task-max normalized AUC of direct score (±std across seeds). Rows include group aggregates (bold) and per-task values.

| Task/Group | AUC(direct score; task-max norm; per step-span) Graph-GA | Unguided optimization | Pointwise guidance | Preference guidance |
|---|---|---|---|---|
| **Similarity** | 0.514±0.019 | 0.365±0.004 | **0.522±0.020** | 0.516±0.016 |
| albuterol_similarity | 0.434±0.017 | 0.315±0.008 | 0.499±0.040 | **0.501±0.031** |
| mestranol_similarity | **0.594±0.033** | 0.416±0.004 | 0.545±0.010 | 0.531±0.005 |
| **MPO** | 0.429±0.010 | 0.370±0.006 | 0.541±0.007 | **0.562±0.014** |
| amlodipine_mpo | 0.395±0.016 | 0.344±0.016 | **0.694±0.007** | 0.676±0.033 |
| fexofenadine_mpo | 0.592±0.031 | 0.580±0.018 | **0.752±0.020** | 0.746±0.010 |
| osimertinib_mpo | 0.666±0.029 | 0.430±0.031 | 0.676±0.015 | **0.684±0.015** |
| perindopril_mpo | 0.409±0.009 | 0.403±0.012 | **0.723±0.013** | 0.691±0.012 |
| ranolazine_mpo | 0.613±0.022 | 0.831±0.008 | **0.857±0.010** | 0.853±0.010 |
| sitagliptin_mpo | **0.113±0.045** | 0.000±0.000 | 0.001±0.001 | 0.020±0.023 |
| zaleplon_mpo | 0.216±0.024 | 0.004±0.008 | 0.083±0.037 | **0.263±0.087** |
| **Rediscovery** | 0.466±0.010 | 0.478±0.005 | **0.635±0.012** | 0.633±0.007 |
| celecoxib_rediscovery | 0.346±0.009 | 0.333±0.009 | 0.517±0.025 | **0.527±0.014** |
| thiothixene_rediscovery | 0.377±0.018 | 0.442±0.010 | **0.595±0.023** | 0.587±0.015 |
| troglitazone_rediscovery | 0.674±0.023 | 0.660±0.004 | **0.793±0.005** | 0.786±0.008 |
| **Hop** | 0.713±0.004 | 0.745±0.003 | 0.784±0.029 | **0.801±0.031** |
| deco_hop | 0.719±0.003 | 0.741±0.005 | 0.781±0.057 | **0.806±0.062** |
| scaffold_hop | 0.707±0.008 | 0.749±0.004 | 0.787±0.013 | **0.796±0.003** |
| **Dock** | 0.266±0.032 | 0.247±0.023 | **0.453±0.031** | 0.449±0.047 |
| drd2_docking | 0.125±0.066 | 0.065±0.017 | **0.308±0.066** | 0.279±0.130 |
| gsk3_beta | 0.320±0.041 | 0.383±0.021 | 0.553±0.040 | **0.590±0.046** |
| jnk3 | 0.352±0.056 | 0.293±0.062 | **0.499±0.054** | 0.478±0.028 |
| **Optimize** | **0.603±0.009** | 0.413±0.006 | 0.508±0.008 | 0.509±0.005 |
| median_1 | **0.575±0.020** | 0.272±0.004 | 0.394±0.011 | 0.407±0.012 |
| median_2 | 0.538±0.010 | 0.635±0.028 | **0.749±0.037** | 0.741±0.019 |
| rdkit_logp | **0.492±0.032** | 0.042±0.002 | 0.064±0.002 | 0.065±0.003 |
| rdkit_qed | 0.659±0.016 | 0.636±0.013 | **0.810±0.010** | 0.808±0.010 |
| sa_tdc | **0.748±0.020** | 0.477±0.007 | 0.522±0.005 | 0.526±0.005 |
| **Isomer** | **0.305±0.021** | 0.000±0.000 | 0.011±0.024 | 0.025±0.053 |
| isomer_c7h8n2o2 | **0.296±0.029** | 0.000±0.000 | 0.000±0.000 | 0.000±0.000 |
| isomer_c9h10n2o2pf2cl | **0.315±0.032** | 0.000±0.000 | 0.022±0.048 | 0.051±0.106 |

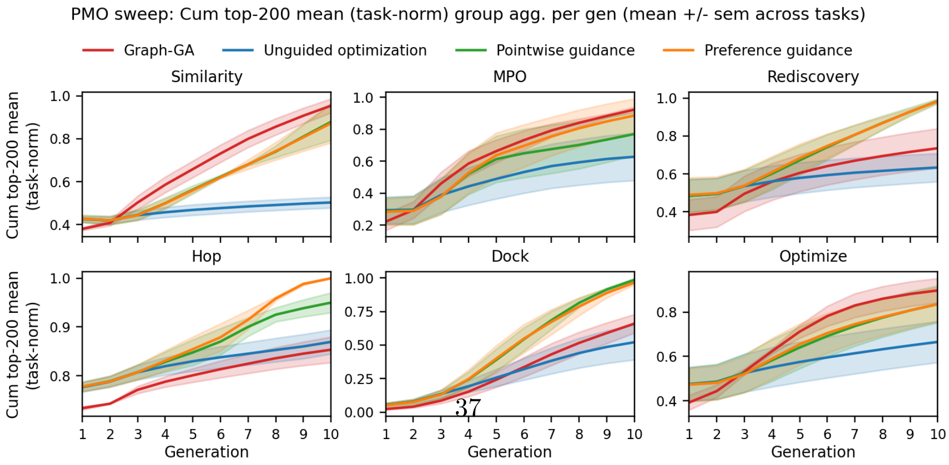




**Supplementary Fig. G3**: PMO sweep grouped aggregates for Cumulative top-200 mean (task-norm). Panels show task-group means across generations for Similarity, MPO, Rediscovery, Hop, Dock, Optimize; shaded bands are +/- sem across tasks. Excluded task(s): rdkit_logp. Groups not displayed: isomer,ungrouped. Values are normalized per task by that task's maximum observed value across all methods and generations before group aggregation.

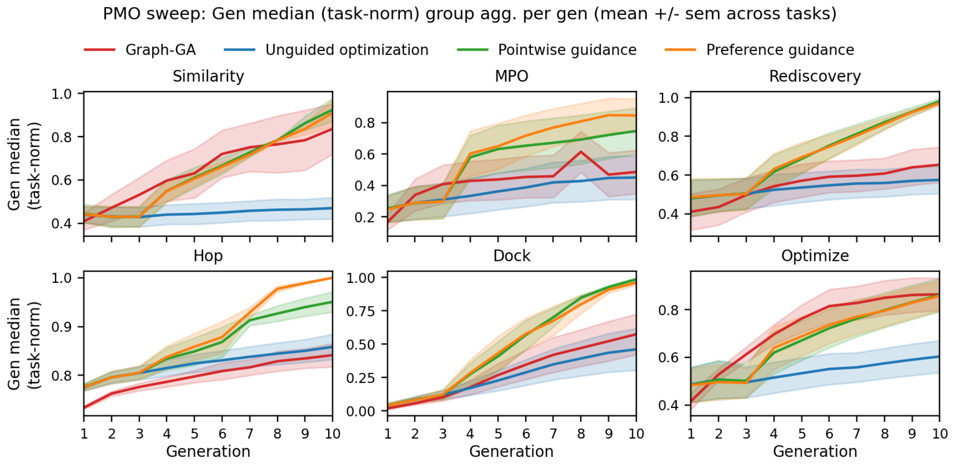


**Supplementary Fig. G4**: PMO sweep grouped aggregates for Gen median (task-norm). Panels show task-group means across generations for Similarity, MPO, Rediscovery, Hop, Dock, Optimize; shaded bands are +/- sem across tasks. Excluded task(s): rdkit_logp. Groups not displayed: isomer,ungrouped. Values are normalized per task by that task's maximum observed value across all methods and generations before group aggregation.

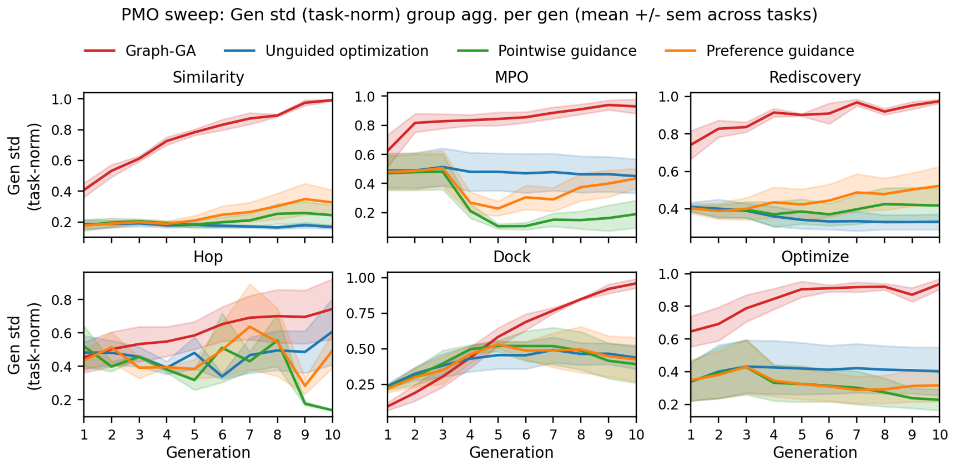


**Supplementary Fig. G5**: PMO sweep grouped aggregates for Gen std (task-norm). Panels show task-group means across generations for Similarity, MPO, Rediscovery, Hop, Dock, Optimize; shaded bands are +/- sem across tasks. Excluded task(s): rdkit_logp. Groups not displayed: isomer,ungrouped. Values are normalized per task by that task's maximum observed value across all methods and generations before group aggregation.

#### G.3 Generation performance on all tasks

The per-task grids below include the `valsartan_smarts` task, which is omitted from the aggregated task-group tables (Table 1, Table G3, and the stepwise variants) because all methods score 0 across seeds.

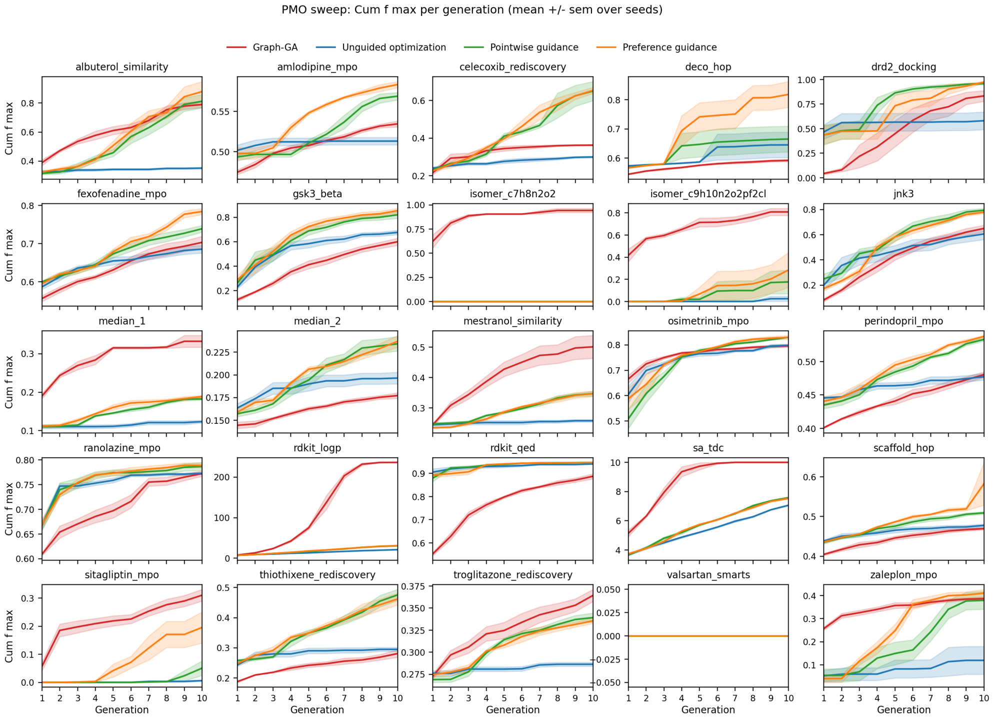


**Supplementary Fig. G6**: PMO sweep generation-wise task-grid figure for Cumulative f max; spread=sem.

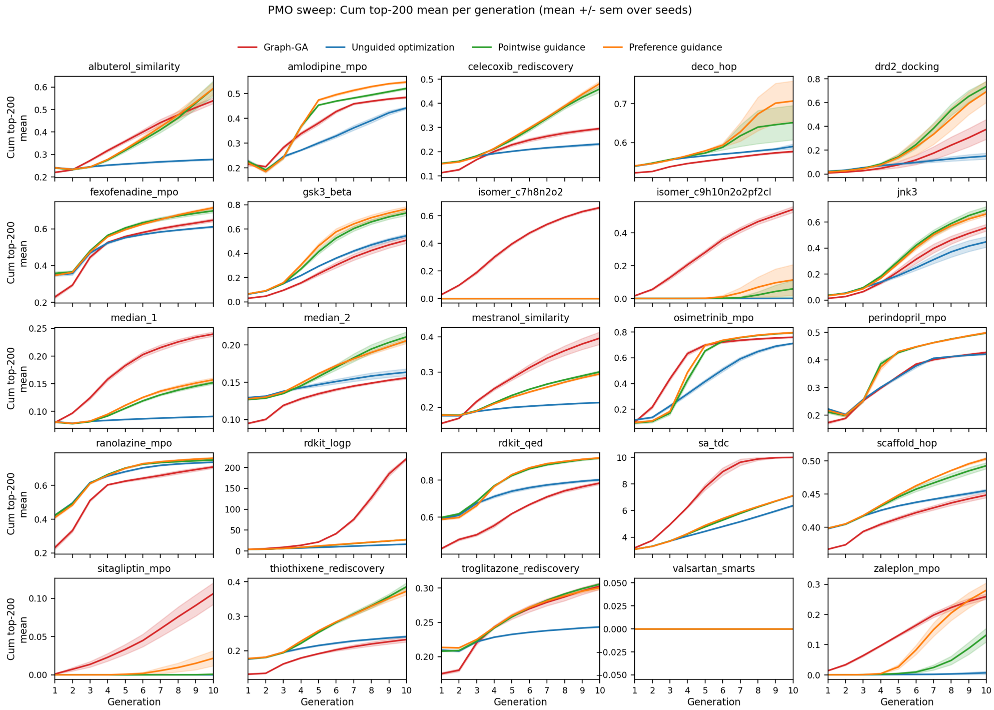


**Supplementary Fig. G7**: PMO sweep generation-wise task-grid figure for Cumulative top-200 mean; spread=sem.

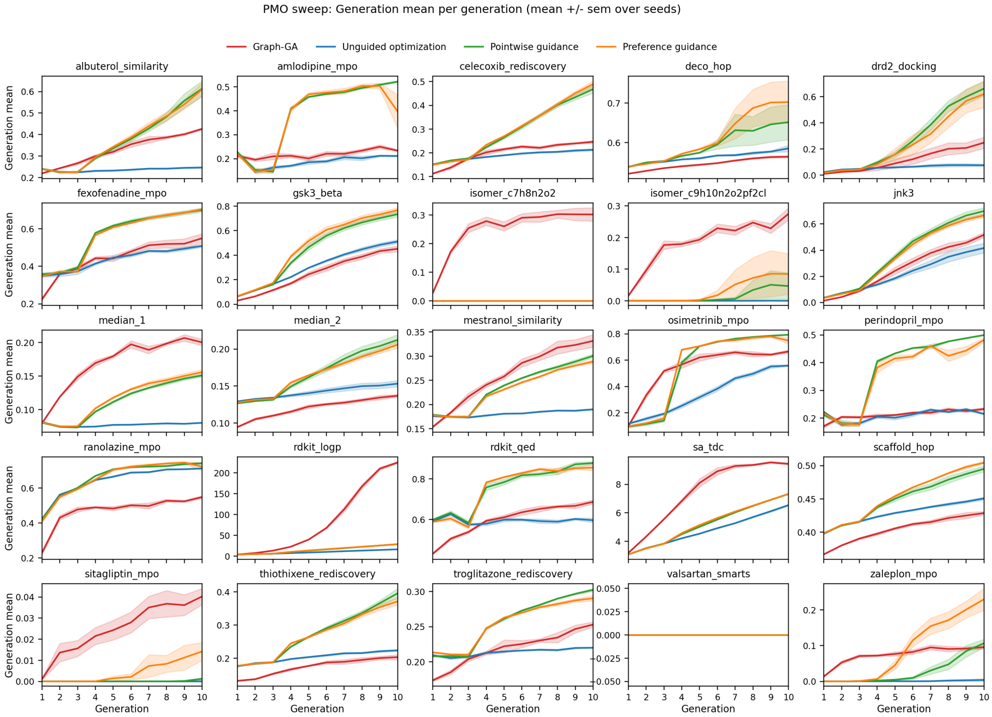


**Supplementary Fig. G8**: PMO sweep generation-wise task-grid figure for Generation mean; spread=sem.

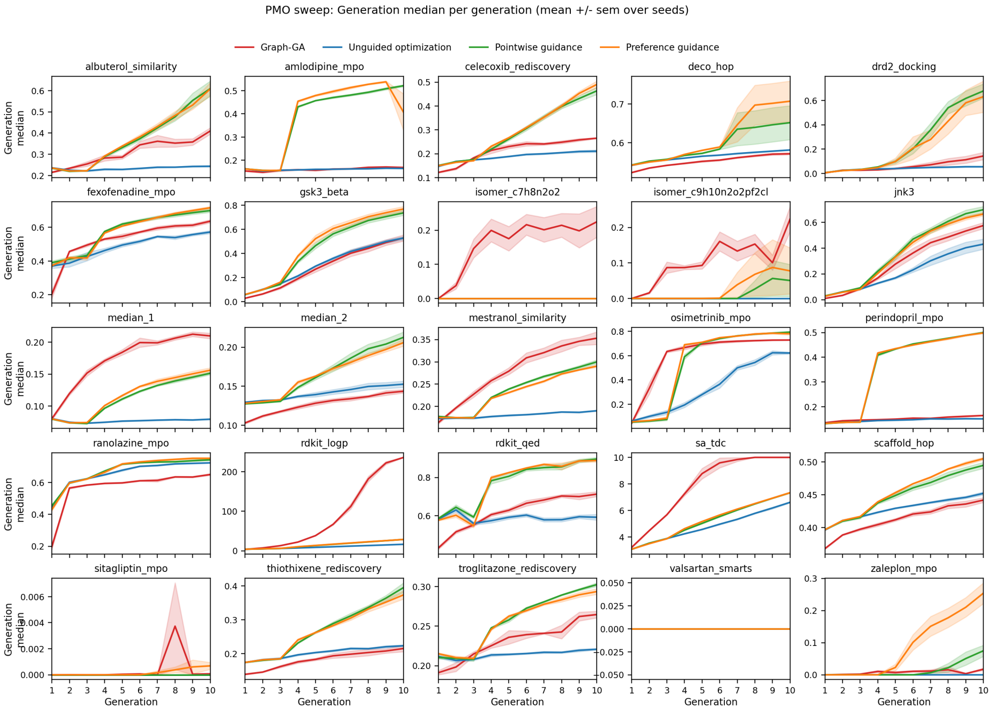


**Supplementary Fig. G9**: PMO sweep generation-wise task-grid figure for Generation median; spread=sem.

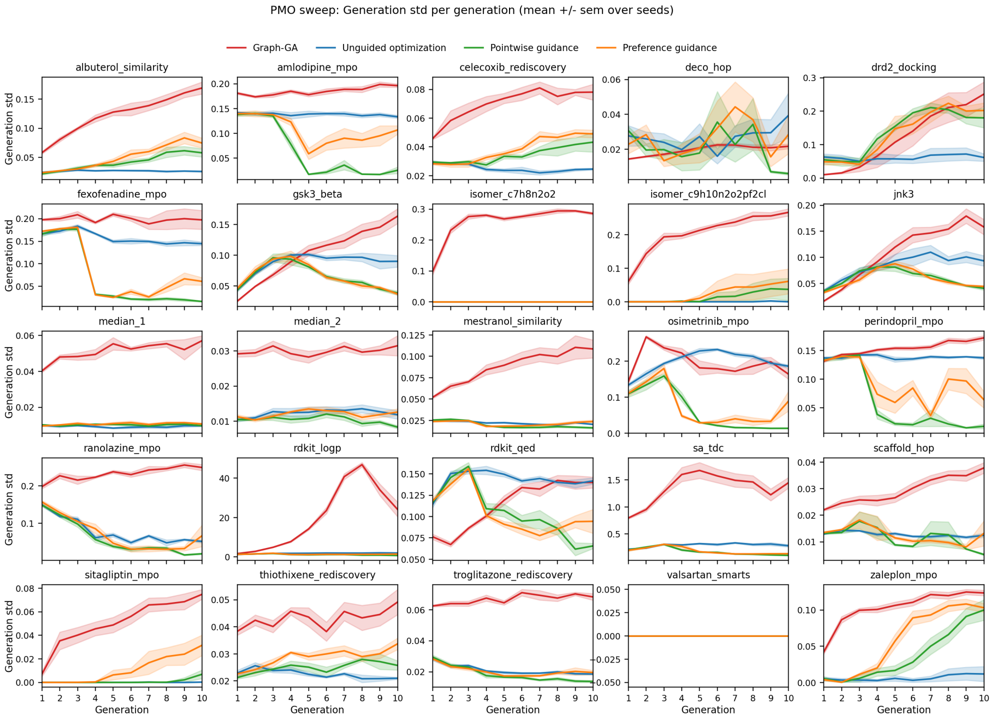


**Supplementary Fig. G10**: PMO sweep generation-wise task-grid figure for Generation std; spread=sem.

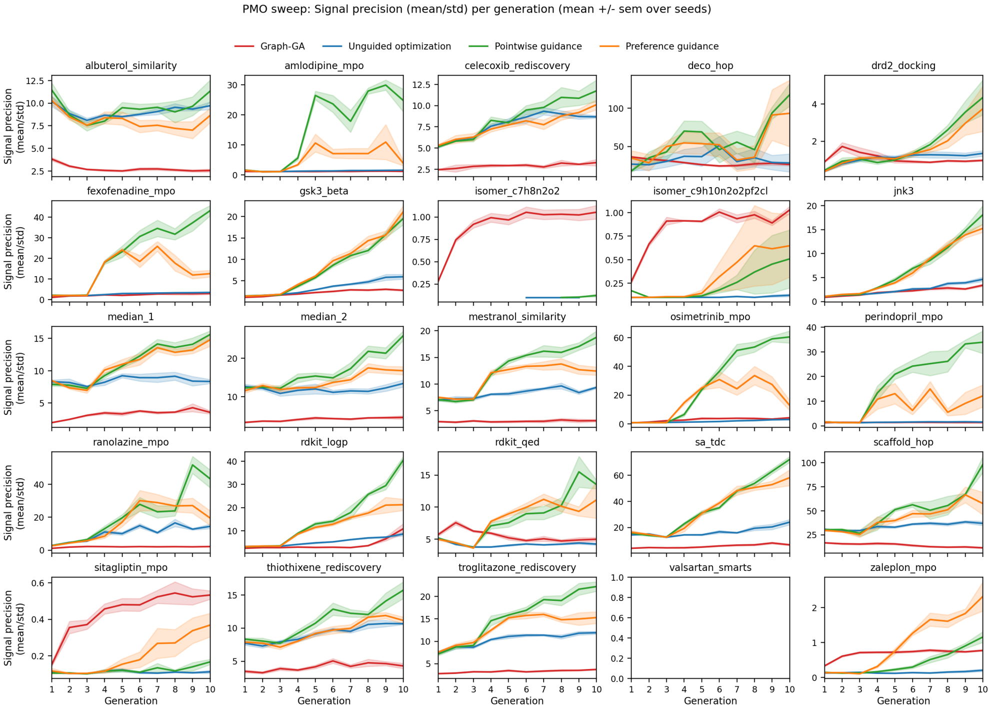


**Supplementary Fig. G11**: PMO sweep generation-wise task-grid figure for Signal precision (mean/std); spread=sem.

##### G.4 Stepwise performance on all tasks

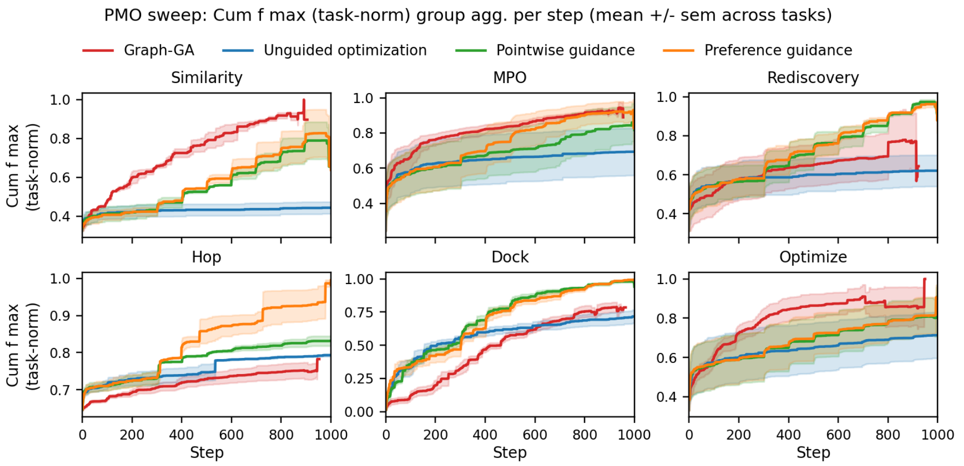


**Supplementary Fig. G12**: PMO sweep stepwise grouped agg. for Cumulative f max (task-norm). Panels show task-group means across optimizer steps for Similarity, MPO, Rediscovery, Hop, Dock, Optimize; shaded bands are +/- sem across tasks. Excluded task(s): rdkit_logp. Groups not displayed: isomer,ungrouped. Values are normalized per task by that task's maximum observed value across all methods and steps before group aggregation.

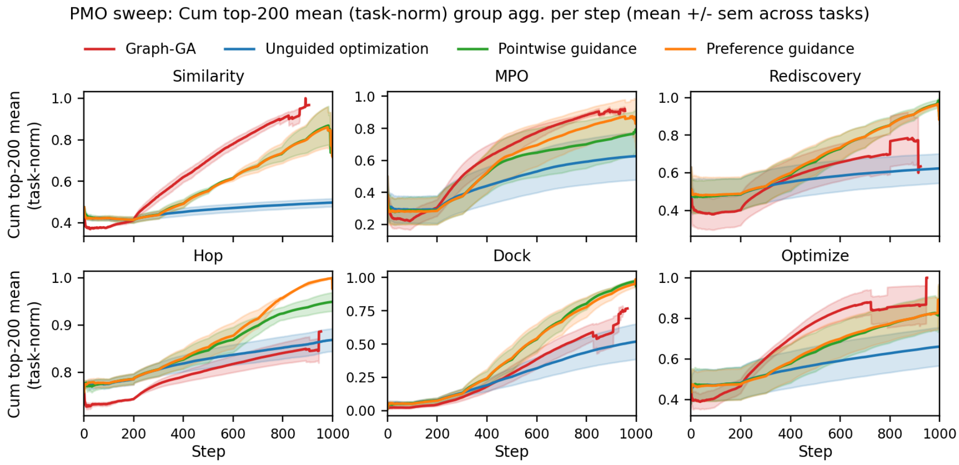


**Supplementary Fig. G13**: PMO sweep stepwise grouped agg. for Cumulative top-200 mean (task-norm). Panels show task-group means across optimizer steps for Similarity, MPO, Rediscovery, Hop, Dock, Optimize; shaded bands are +/- sem across tasks. Excluded task(s): rdkit_logp. Groups not displayed: isomer,ungrouped. Values are normalized per task by that task's maximum observed value across all methods and steps before group aggregation.

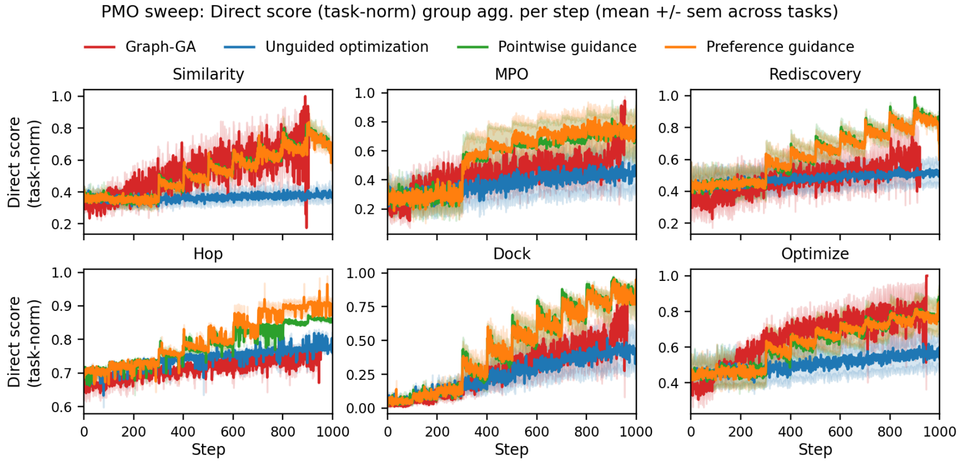


**Supplementary Fig. G14**: PMO sweep stepwise grouped agg. for Direct score (task-norm). Panels show task-group means across optimizer steps for Similarity, MPO, Rediscovery, Hop, Dock, Optimize; shaded bands are +/- sem across tasks. Excluded task(s): rdkit_logp. Groups not displayed: isomer,ungrouped. Values are normalized per task by that task's maximum observed value across all methods and steps before group aggregation.

#### G.5 Diversity of generated samples

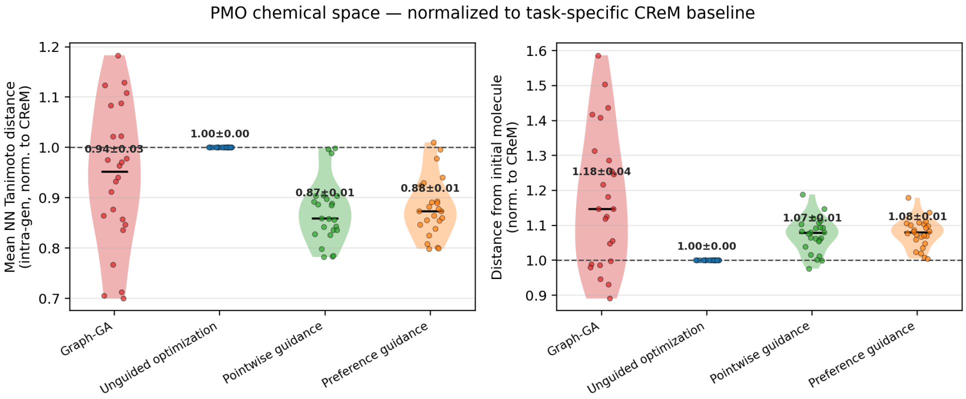


**Supplementary Fig. G15**: **Chemical-space diversity of PMO runs, normalized to the CReM baseline.** Violin plots show the distribution of per-task diversity scores across the 24 PMO benchmark tasks (`rdkit_logp` excluded), with individual task means overlaid as scatter points and the median indicated by a horizontal bar. Each value is expressed as a ratio to the task-specific `CReM` mean (dashed line at 1), so that values above 1 indicate greater diversity than the `CReM` baseline and values below 1 indicate less. **(Left)** Mean nearest-neighbour Tanimoto distance within a generation (intra-generation diversity) across 10 generations: higher values reflect a more chemically diverse set of molecules proposed in a single generation. **(Right)** Mean Tanimoto distance of current-generation molecules from the initial molecule (exploration from the starting point): higher values indicate that the optimizer has moved further from the seed structure. Annotations show mean ± SEM across the 24 tasks.

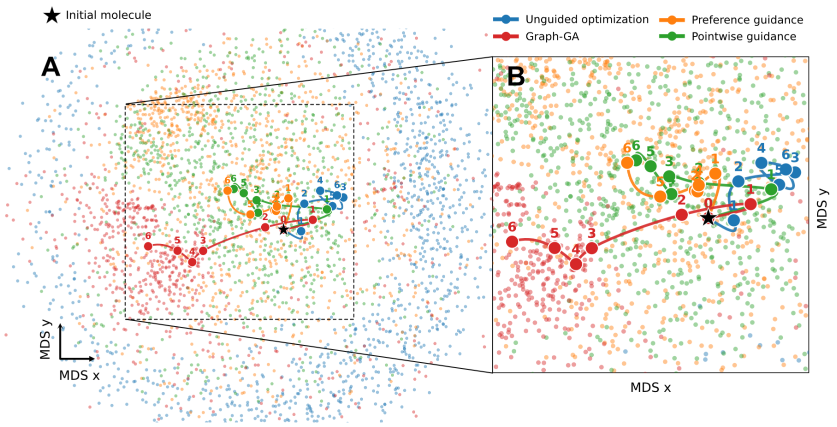


**Supplementary Fig. G16**: **A/B**: All sampled molecules from a single seed of the GPCR ligand rediscovery tasks viewed along the fitness axis, with inset **B** showing a close-up of the explored region and each of the different methods for optimization. The unguided optimization in blue covers the largest area of in the dimensionality-reduced chemical space. The trajectory points are computed from the mean MDS x and MDS y in each generation for each method.
